# Comprehensive characterization of V(D)J recombination from long-read transcriptomic data with VDJcraft

**DOI:** 10.64898/2026.04.01.715879

**Authors:** Kaili Hu, Alexander Rosenberg, Yuwei Song, Chia-Hsuan Fan, Zishan Peng, Min Gao, Zechen Chong

## Abstract

V(D)J recombination generates antigen receptor diversity in developing B and T cells. Long-read transcriptome technologies (e.g., PacBio Iso-Seq, Nanopore RNA/cDNA) capture full-length transcripts and thus resolve V(D)J events more accurately than short-read platforms. However, existing short-read tools are not applicable to or optimized for long-read data. We developed VDJcraft, the first integrated pipeline designed for V(D)J recombination analysis using long-read transcriptome sequencing data. The workflow uses a two-pass alignment strategy: global alignment to the GENCODE reference with minimap2, followed by local realignment and annotation using the international ImMunoGeneTics information system (IMGT). A customized module enhances D-gene detection sensitivity and positional precision. Sequencing errors are reduced through consensus-based correction toward the predominant subclass. Antigen-binding regions are annotated using IMGT-defined motifs to characterize CDRs and binding site composition. VDJcraft was validated on simulated and Human Genome Structural Variation Consortium (HGSVC) datasets and applied to disease datasets. It accurately recovered full-length V(D)J-C sequences and outperformed existing methods in gene detection and recombination accuracy. Long-read calls also showed significantly higher concordance with high-confidence short-read calls (Mann-Whitney U test, p = 1.55 × 10⁻^4^). Additionally, we identified 31 putative novel gene subclasses absent from the IMGT database from HGSVC datasets. Analyses of longitudinal blood samples from a COVID-19 patient revealed distinct V(D)J recombination patterns and segment enrichment, characterized by increased IGHV1-2 usage, enrichment of the IGHV3-7/IGHD6-9/IGHJ5_02 rearranged clonotype, and a transient peak in IgG2 levels at day 4 followed by a gradual return to baseline. In conclusion, VDJcraft provides a robust framework for long-read V(D)J characterization and enables the discovery of disease-associated immune signatures.

## Introduction

The immune system is an intricate network comprising lymphoid organs, various cell types, humoral factors, and cytokines^1^. The effective functioning of this system is vital for maintaining health, as its dysfunction can lead to immunodeficiency disorders, resulting in life-threatening infections^2^ and tumor development^3^, while an overactive immune system can cause allergic reactions and autoimmune diseases^4,5^. A fundamental process underlying the adaptive immune system is V(D)J recombination, which plays a pivotal role in generating the vast diversity of antigen receptors necessary for immune defense^6,7^. This genetic recombination process occurs during the development of B- and T-lymphocytes, enabling the production of highly diverse immune receptor repertoires^3,8^. Through this mechanism, immunoglobulins and T-cell receptors acquire the ability to recognize and bind specific antigens, providing a tailored immune response to pathogens and other foreign substances. Despite decades of research, the entire landscape of V(D)J recombination remains not fully understood, hindering our ability to fully elucidate the mechanisms shaping the immune repertoire^6,8^. This is largely attributed to the complexity and variability of the process.

V(D)J recombination has been extensively studied over the past decades. Conventional methods such as PCR^9^, Quantitative PCR^10^, RT-PCR^10,11^, Southern blot hybridization^12^, and Sanger sequencing^13^ have played a significant role in analyzing this process. While these techniques have provided valuable insights into the mechanisms of V(D)J recombination, they are inherently limited due to constraints in product length, primer bias, and coverage. As a result, these approaches can only capture a small subset of the immune repertoire and lack the ability to provide a comprehensive view of its diversity^14^. With advancements in sequencing technologies, high-throughput methods, including next-generation and third-generation sequencing, have revolutionized our ability to study V(D)J recombination, offering a more complete and comprehensive characterization of immune repertoire diversity. These technologies have paved the way for the development of bioinformatic tools, which enable the detailed identification and characterization of immune receptor repertoires. By leveraging these technologies, researchers continue to gain deeper insights into the intricate processes underlying the immune diversity and function^15–17^.

With the rapid advancement of next-generation sequencing (NGS), several bioinformatics tools have been developed to analyze V(D)J recombination. Tools such as IMGT/HighV-QUEST^18,19^, developed by IMGT, the international ImMunoGeneTics information system^20^, provide standardized analysis of V(D)J identifications. However, they are web-based, which limits the number of sequences that can be analyzed per batch. Other tools, such as TRUST4^21^, LymAnalyzer^22^ have improved the efficiency and functionality of V(D)J recombination analysis, providing deeper insights into immune repertoire. These tools are capable of detecting V(D)J rearrangements from short-read sequencing data; however, they are inherently optimized for short reads. As a result, they face significant challenges when applied to long genes, such as V genes, where short-read lengths can span only partial segments of the locus. Such partial coverage can lead to incomplete alignments and mapping ambiguities, particularly in regions with high sequence similarity or repetitive elements. These limitations can increase error rates^23,24^, hinder accurate gene segment assignment, and result in the loss of critical sequence information^25^, ultimately limiting the accuracy and comprehensiveness of immune repertoire characterization. Notably, among the currently available short-read-based tools, TRUST4, outperforms others with its advanced algorithm for immune repertoire analysis from both bulk RNA-seq and single-cell RNA-seq (scRNA-seq) data, excelling in precision and speed. Despite these advancements, TRUST4 has limitations in sensitivity and is prone to potential errors in its *de novo* assembly process, which can compromise its ability to accurately characterize V(D)J recombination. Consequently, current tools remain insufficiently accurate and comprehensive to fully characterize the complete landscape of V(D)J recombination and the immune repertoire.

The emergence of third-generation sequencing technologies has transformed genomic analysis through long-read sequencing platforms such as PacBio and Oxford Nanopore (ONT). These technologies generate full-length sequencing reads, offering superior accuracy and completeness compared to traditional short-read methods^26,27^. Long-read sequencing has the potential to capture the entire V(D)J rearrangements, allowing for more precise and comprehensive analysis of the immune repertoire **(Supplementary Figure 1).** To fully leverage the advantages of long-read technologies, it is imperative to develop robust and efficient bioinformatics tools specifically designed for processing and analyzing such data. These tools will facilitate accurate exploration and characterization of immune repertoires, thereby enhancing downstream analyses and advancing our understanding of immune system diversity and function.

Here we present VDJcraft (**Figure 1**), a pipeline designed for comprehensive analysis of V(D)J recombination in the immune repertoire using PacBio and ONT (Oxford Nanopore Technology) long-read transcriptomic data. VDJcraft enables precise gene identification, including V, D, J, and C genes, as well as CDR1, CDR2, and CDR3 regions, while also capturing somatic hypermutation and class-switch events. The pipeline aligns long-read transcriptomic data to the human genome and extracts candidate reads mapping to V, D, J, and C genes based on their genomic location. It then performs local alignment of these candidate reads to the IMGT database using BLAST with customized parameters, ensuring accurate identification of V, D, J, and C genes through stringent filtering criteria (Methods). CDRs are determined by targeting conserved amino acid positions or traditional IMGT-defined locations and consensus sequences. To enhance identification accuracy, VDJcraft incorporates error corrections during processing. VDJcraft demonstrates superior performance and exceptional precision, positioning it as a highly effective and reliable tool for characterizing V(D)J recombination using long-read sequencing technologies. Its advanced capabilities address critical limitations of existing approaches, providing a comprehensive solution for analyzing the complex dynamics of the immune system.

**Figure 1.**
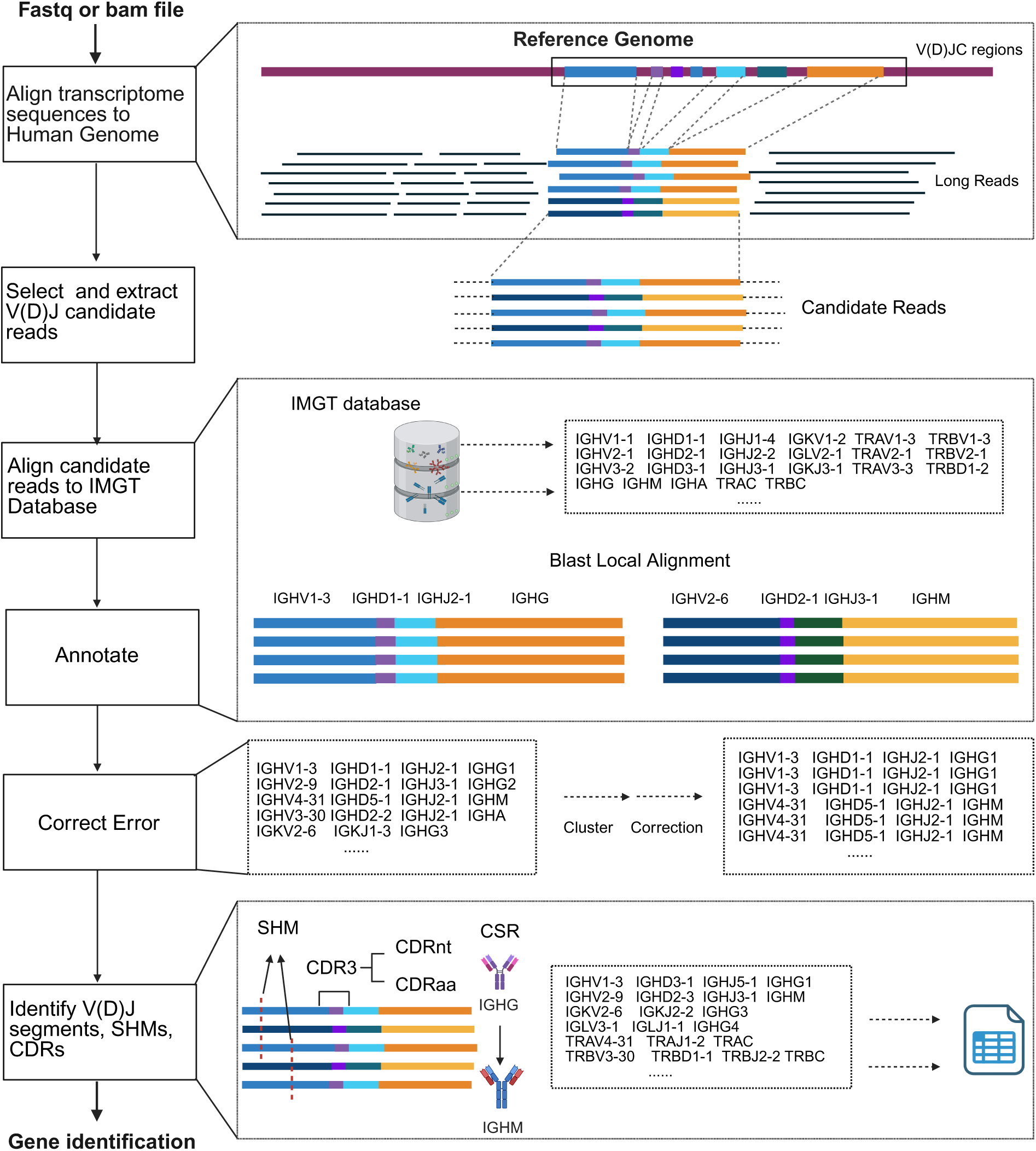
The workflow of VDJcraft. VDJcraft, an integrated pipeline for analyzing V(D)J recombination using long-read transcriptomic data. Taking long-read transcriptome sequences as input, a two-pass alignment strategy is employed to accurately capture full-length V, D, and J gene segments. Reads are initially aligned to the reference genome (e.g. GENCODE) using minimap2, followed by local realignment and annotation by aligning to the IMGT database. A customized alignment module is implemented to enhance D gene detection sensitivity and positional accuracy. Sequences are corrected to the predominant subclass by employing a consensus-based strategy to mitigate sequencing errors. Antigen-binding domains are explored using IMGT-defined consensus motifs to delineate CDRs and infer binding site composition for repertoire diversity analysis.

## Methods

### VDJcraft workflow

#### Overview of VDJcraft

VDJcraft is a pipeline for comprehensively characterization of V(D)J recombination in long-read transcriptomes. The workflow consists of the following functional steps: First, VDJcraft aligns long-read transcriptomic data to the latest version of the genome reference from GENCODE using Minimap2. At the same time, it generates BED files containing the genomic coordinates of V, D, J, and C genes. These BED files are then used to target the V(D)J regions on the long reads and extract V(D)J candidate reads. Next, the candidate reads are aligned to the IMGT V(D)J database using BLAST. The IMGT database meticulously maintains sequences for V, D, J, and constant genes^20^ and serves as a primary reference for annotating BCR and TCR sequences in widely used tools such as TRUST4^21^. This step produces a list of V, D, J, and C genes with their corresponding sequences and annotations. Finally, VDJcraft employs the IMGT CDRs consensus sequences to identify CDR1 and CDR2 within the germline V gene sequences, and generates CDR3 by joining the V, D and J segments. The CDRs are then translated to obtain the corresponding amino acid sequences. VDJcraft is implemented as a Python program for the identification of V(D)J rearrangements in third generation sequencing data. It can identify full-length V(D)J gene sequences with some achieving up to 100% sequence identity to entries in the IMGT database.

#### Candidate reads extraction

VDJcraft identifies candidate TCR and BCR reads from raw sequencing data (FAFTQ/FASTA format) or from alignment files (BAM format) generated by long-read aligners such as minimap2^28^. When raw FASTQ or FASTA files are provided, VDJcraft first aligns the reads to the selected reference genome to generate a corresponding BAM file. Candidate reads are then extracted based on V, J, and C gene locus BED files that are bundled with the pipeline. Multiple genome references are supported for alignment, including Telomere-to-Telomere (T2T), GRCh38, GRCh37, and GRCm39, each with built-in BED files specifying V, J, and C loci. When a BAM file is provided as input, VDJcraft directly extracts the aligned reads as the candidate read set. In this case, the use must specify the reference genome used for the original alignment, such as “-M” when supplying GRCm39-aligned reads.

#### Customization for gene identification and annotation

VDJcraft implements a rigorous and systematic approach for identifying the V, J, and C genes of T-cell receptors (TCRs) and B-cell receptors (BCRs). The process begins by analyzing BLAST local alignment results and selecting the highest-scoring alignment for each read after applying quality filters. Specifically, alignments with an identification percentage below 90% (the default threshold, adjustable as needed) were excluded, and V genes with a score below 350 (the default threshold, adjustable as needed) were additionally filtered to ensure accuracy. For each read, the top-ranking alignments that meet or exceed this threshold are retained. When multiple alignments have identical scores, VDJcraft selects the match with the greatest identification percentage and aligned length. After filtering, VDJcraft compiles a comprehensive list of the best-scoring matches and maps them to gene annotations from the IMGT database. This procedure ultimately assigns the corresponding V, J, and C genes for each long read, providing precise gene characterization essential for downstream analyses.

Accurate identification of D genes remains challenging due to their short length and extensive sequence diversity. To address this, VDJcraft incorporates a targeted alignment strategy that enhances the precision of D gene detection and integrates seamlessly with the V, J, and C gene analysis. The workflow first identifies the V and J genes within each read and determines their genomic positions. Using these coordinates, VDJcraft extracts the intervening sequence between the V and J segments. To increase the likelihood of capturing D genes, an additional 20 base pair (bp) are included on both sides of the V-J interval. These extracted sequences are then aligned against the IMGT D gene database to identify candidate D genes. This approach enables accurate detection of short D genes that may otherwise be missed during full-length sequence alignment. To further improve specificity, VDJcraft filters out low-confidence D gene matches by retaining only alignments with an identification percentage above 80% (default threshold, adjusted as needed). For each read, the highest-scoring D gene accurate assignment. These annotations are then incorporated into the comprehensive VJC identification list while preserving the corresponding read names. Additionally, VDJcraft can recover complete VDJ sequences for reads in which short D genes are not captured through database alignment by leveraging previously determined V and J positions. This systematic approach not only improves D gene detection accuracy but also ensures their robust integration into downstream analyses.

#### CDRs extraction

VDJcraft analyzes candidate sequences from the international ImmunoGeneTics (IMGT) database to discern V, J, and C genes. In addition to gene-segment identity, IMGT provides annotations of complementary determining regions (CDRs) based on positional definitions. Specifically, VDJcraft determines the initiation site of CDR3 at the conserved cysteine (C) located at the end of V gene within the YYC motif. If this motif is absent, the CDR3 start position is defined as the 104th amino acid, following IMGT conventions. The termination site of CDR3 is identified as the tryptophan (W) or phenylalanine (F) within the W/FGxG motif of the J gene, consistent with the definition used by TRUST4^21^. After VDJcraft assigns the V and J genes, it computes CDR3 coordinates based on these IMGT rules. IMGT defines CDR1 (positions 27–38) and CDR2 (positions 56–65) for immunoglobulin (IG) and T-cell receptor (TR) genes, and anchors these regions with conserved amino acids, a cysteine at position 23 and a tryptophan at position 41^20^. VDJcraft identifies and extracts CDR sequences by applying the IMGT-defined positional boundaries together with the consensus sequence of each V gene, ensuring accurate and standardized delineation of CDR1 and CDR2 across allelic diversity.

#### Error correction

To address the higher error rates associated with certain long-read sequencing platforms, such as Oxford Nanopore Technology, VDJcraft incorporates a robust error correction module designed to enhance the accuracy of V(D)J recombination identification and streamline downstream analyses. This module identifies and corrects confidently supported sequencing errors, including those that are supported by a single or double-read alignments. Operating within a defined 50 base pair (bp) window, the module groups sequencing errors into subclusters based on the alignment start positions of sequencing reads. Within each subcluster, VDJcraft prioritizes the most frequently observed V, D, J, and C gene annotation combinations for correction. Minor deviations—defined as cases in which only one of the V, D, J, or C annotations differs from the dominant combination—are corrected to match the dominant gene assignment. This targeted correction process specifically addresses inaccuracies arising from sequencing-base errors or from gene identification supported by limited read evidence, thereby improving the accuracy of immune receptor characterization,

Formally:

· Each recombined sequence ‘s’ has gene annotations *G(s),*

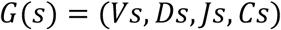

where 𝑉𝑠, 𝐷𝑠, 𝐽𝑠, 𝑎𝑛𝑑 𝐶𝑠 represent V, D, J, and C gene segments, respectively.

• Let the dominant annotation combination in the dataset be

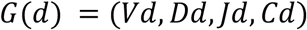

where *Vd*, *Dd*, *Jd*, and *Cd* represent the dominant gene of the recombination, respectively.

· Define the deviation count between a sequence and the dominant combination as

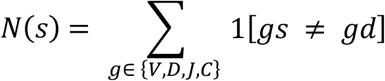

where the number of 𝑉𝑠, 𝐷𝑠, 𝐽𝑠, 𝑎𝑛𝑑 𝐶𝑠 that deviates from the dominant *Vd*, *Dd*, *Jd*, and *Cd* within a 50-bp window is counted.

· Error correction rule:

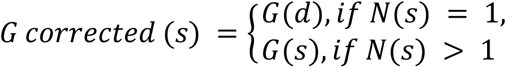

If the deviation count equals 1, the gene segment is corrected to the dominant gene segment; if the deviation count is greater than 1, the gene segment(s) are retained as originally assigned.

### Benchmark on simulation datasets

#### Simulation dataset generation

Simulated immunoglobulin and T cell receptor sequences with ground truth were generated for benchmarking. For immunoglobulin, 200 IG heavy-chain sequences, 200 IG (𝜅, 𝜆) light-chain sequences were randomly selected as reference sequences. For T cell receptors, 100 TRA sequences and 100 TRB sequences were selected. These sequences were recombined in a randomized fashion, and half were converted to reverse-complement sequences. Mutations were randomly introduced at a rate of 1x10^-4^. Furthermore, to model differences in transcript abundance across different sequencing platforms, expression levels were simulated at high (40x), medium (20x), and low (10x) coverage. PacBio-like reads were simulated from the modified genome using pbsim (v1.0.3) with the options “--length-mean 2000 --length-min 1000 --length-max 3500 --model_qc CCS --accuracy-max 1.00 --accuracy-min 0.98”. Illumina-like reads were simulated using Art-illumina^29^ with the options “-ss HS25 -sam -p -l 150 -m 200 -s 10 -c 20x”.

#### Gene discovery in simulated datasets

VDJcraft was applied to all simulated datasets using default parameters. The GRCh38 human reference genome was used for alignment. TRUST4 and LymAnalyzer were applied to the simulated Illumine datasets. Gene detections were compared against ground truth to assess accuracy. The reported V(D)J gene identifications and complete recombination events from VDJcraft, TRUST4, and LymAnalyzer were compared to calculate recall, precision, and F1 scores.

F1 score:

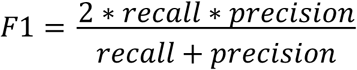

Commands used:

· TRUST4:

*run-trust4 -f IMGT.fa --ref IMGT.fa -1 R1_001.fastq.gz -2 R2_001.fastq.gz*

· LymAnalyzer (IGH):

*java -jar -Xmx4g LymAnalyzer_cmd_1.2.2.jar simu.fastq lymout IGH hs lymigh No No 10 GRCh38.fa*

#### Simulation of ultra-long nanopore reads and cross-platform comparison

Nanopore-like reads were simulated using Badread (v0.2.0) ^30^with the options *“--quantity 20x – error_model random –qscore_model ideal –glitches 0,0,0 –junk_reads 0 –random_reads 0 – chimeras 0 –identity 30,3 –length 40000,20000 ”*.

VDJcraft was employed to both PacBio-like and Nanopore-like reads at 20x coverage. Identified V(D)J gene segments and recombination events were compared to the ground truth to calculate recall, precision, and F1 scores for each platform.

### Benchmarking and evaluation of VDJcraft on HGSVC Data

#### Gene discovery and comparison between VDJcraft and TRUST4

Raw reads from 12 PacBio Iso-Seq datasets (HG02106, HG01457, HG00268, HG02666, HG03248, HG03807, HG04217, NA18989, NA19317, NA19331, NA19347, and NA19384) were downloaded from the Human Genome Structural Variation Consortium (HGSVC)^31^, along with corresponding high-coverage short-read (Illumina) RNAseq data. VDJcraft was applied to the PacBio Iso-Seq datasets, whileas TRUST4 was applied to the short-read data. Individual gene segments and VJ recombination identified by both tools were compared to calculate consistency. Comparisons were performed before and after applying a TRUST4 threshold of > 0.1% clonotype frequency, based on support reads. CDR3 amino acid sequences detected by VDJcraft and TRUST4 were also compared, with a focus on top 50 mostly enriched TRUST4 results.

#### Somatic hypermutation detection and novel event discovery

VDJcraft includes a somatic hypermutation (SHM) detection module that leverages inexact local alignment to IMGT reference gene segments under relaxed identity thresholds. Sequences with <85% identity to IMGT entries were classified as containing SHMs. Within each HGSVC sample, unique sequences were identified and enumerated, and those supported by only one read were excluded to minimize false positives. Inexact alignments to IMGT entries supported by more than two reads were classified into corresponding immune gene categories but lacked exact IMGT matches, indicating potential novel hypermutated events.

### Application and validation of VDJcraft in FLAIRR-Seq healthy and COVID-19 samples

#### Validation of V(D)J recombination detection using healthy cohort data

VDJcraft was utilized to analyze real data comprising the IgG repertoires of ten healthy individuals generated by FLAIRR-Seq^32^ that are publicly available on the Sequence Read Archive (BioProject PRNJA922682). FLAIRR-Seq is a method for profiling near full-length antibody heavy chain repertoires, enabling the generation of highly accurate antibody heavy chain transcripts using PacBio long reads. By leveraging VDJcraft, V(D)JC genes were accurately identified and annotated. IGHV subclass enrichment patterns and IgG subclasses across different IGHV genes were subsequently analyzed, allowing direct comparison and cross-validation with results previously reported in the FLAIRR-Seq original work^32^.

#### Application of VDJcraft for profiling immune infiltration in COVID-19

IgG heavy-chain FLAIRR-seq datasets from whole-blood samples of a hospitalized COVID-19 patient, collected longitudinally on days 1, 4, 8, and 13 after hospital admission, were analyzed using VDJcraft. The tool identified dynamic IGHV usage across the recovery period, detected IGHG class-switch recombination patterns, and captured shifts in V(D)J recombination frequencies. Furthermore, CDR3 motif prediction using Memesuite^33^ and downstream functional characterization were conducted, offering insights into transient immune repertoire changes during the patient’s recovery from COVID-19.

## Data availability

HGSVC PacBio Iso-seq data are available at the HGSVC data portal: https://ftp.1000genomes.ebi.ac.uk/vol1/ftp/data_collections/HGSVC3/working/20210728_PacBio_Iso-Seq_JAX/;

Illumina RNAseq data for HGSVC samples were generated at the Jackson laboratory for Genomic Medicine on behalf of the Human Genome Structural Variation Consortium. https://ftp.1000genomes.ebi.ac.uk/vol1/ftp/data_collections/HGSVC3/working/2021_RNAseq_JAX/.

FLAIRR-seq are available in the Sequence Read Archive under BioProject PRNJA922682.

## Code availability

VDJcraft is publicly available under the MIT License (https://github.com/kailihu-uab/VDJcraft). Version 1.1.1 was used for all V(D)J recombination detection and benchmarking presented in this study. Key custom Python scripts used in the manuscript are available at https://github.com/kailihu-uab/VDJanalysis.

## Results

### Benchmark on simulated datasets

To benchmark the accuracy of V(D)J recombination detection, we compared VDJcraft with two other bioinformatic tools, TRUST4 and LymAnalyzer, on a simulation dataset. TRUST4 and LymAnalyzer could not be successfully applied to the HiFi long-read data from the HGSVC sample HG00268, indicating that these tools are incompatible with long-read data. Therefore, we generated simulated long reads for VDJcraft and simulated short reads for TRUST4 and LymAnalyzer for comparison.

Under the default settings, we first analyzed individual gene detection using VDJcraft, TRUST4 and LymAnalyzer on simulated datasets (**Table 1**), VDJcraft demonstrated superior recall (sensitivity) and precision for identifying IGHV and IGHD genes compared to TRUST4 and LymAnalyzer, while all three tools achieved high accuracy in detecting IGHJ genes. For light chain genes, VDJcraft and TRUST4 consistently outperformed LymAnalyzer in both recall and precision across IGK(L)V and IGK(L)J genes. Notably, for IGK(L)V genes, VDJcraft exhibited the highest accuracy, surpassing both TRUST4 and LymAnalyzer in recall and precision. In the case of T-cell receptor genes (TRA and TRB), VDJcraft and TRUST4 exhibited higher recall and precision than LymAnalyzer, with consistently high precision across both tools; however, VDJcraft achieved greater recall than TRUST4. Overall, VDJcraft exhibited superior accuracy in detecting individual V, D, and J genes across both heavy and light chains, for both T cells and B cells. This advantage was particularly pronounced in the detection of long V gene segments. VDJcraft achieved recalls of 1.00 for IGHV, 0.98 for IGK(L)V, 0.98 for TRAV and 1.00 for TRBV, with precision values ranging from 0.90 to 1.00. In contrast, TRUST4 showed reduced recall for IGHV (0.92), while LymAnalyzer achieved moderate recall (0.90) but poor precision (0.50). Consequently, VDJcraft demonstrated a markedly higher F1 score of 0.95 for V gene detection. This performance substantially surpassed the F1 scores of TRUST4 (0.87) and LymAnalyzer (0.64), highlighting the efficacy and accuracy of VDJcraft in capturing V gene diversity (**Table1**). These results underscore VDJcraft’s robust performance across diverse gene classes and immune receptor types, demonstrating its strong potential as a highly accurate tool for long-read-based V(D)J recombination analysis.

**Table 1.** Performance comparison of individual V(D)J gene segment identification on simulated datasets.

| Genes | VDJcraft |  |  | TRUST4 |  |  | LymAnalyzer |  |  |
| --- | --- | --- | --- | --- | --- | --- | --- | --- | --- |
|  | Recall | Precision | F1 score | Recall | Precision | F1 score | Recall | Precision | F1 score |
| IGHV | **1.00** | **0.90** | **0.95** | 0.92 | 0.82 | 0.87 | 0.90 | 0.50 | 0.64 |
| IGHD | **1.00** | **0.94** | **0.97** | 0.87 | 0.77 | 0.81 | 0.87 | 0.87 | 0.87 |
| IGHJ | **1.00** | **1.00** | **1.00** | **1.00** | **1.00** | **1.00** | **1.00** | 0.91 | 0.95 |
| IGK(L)V | **0.98** | **0.97** | **0.98** | 0.74 | 0.90 | 0.81 | 0.79 | 0.55 | 0.65 |
| IGK(L)J | 0.95 | **1.00** | **0.97** | 0.95 | **1.00** | **0.97** | **1.00** | 0.95 | **0.97** |
| TRAV | **0.98** | **1.00** | **0.99** | 0.82 | 0.93 | 0.82 | 0.54 | 0.81 | 0.65 |
| TRAJ | **1.00** | **1.00** | **1.00** | **1.00** | **1.00** | **1.00** | 0.67 | 0.67 | 0.67 |
| TRBV | **1.00** | **0.97** | **0.99** | 0.93 | 0.90 | 0.92 | 0.70 | 0.42 | 0.53 |
| TRBD | **1.00** | **1.00** | **1.00** | **1.00** | **1.00** | **1.00** | **1.00** | **1.00** | **1.00** |
| TRBJ | **1.00** | **1.00** | **1.00** | 0.94 | **1.00** | 0.97 | 0.94 | **1.00** | 0.97 |

Results comparing the accuracy of individual V(D)J gene segment detection by VDJcraft, TRUST4, and LymAnalyzer on simulated dataset. Recall, Precision, and F1 scores are reported for each gene segment. Bold values indicate the highest Recall, Precision, or F1 score among VDJcraft, TRUST4, and LymAnalyzer for each gene segment. Bold font values indicate the highest F1 score in the same gene category of each row.

To assess the full length of V(D)J recombination representing clonotypes, we analyzed VDJ rearrangements for B- and T-cell heavy chains and VJ rearrangements for their light chains. As shown in **Figure 2**, for V(D)J recombination detection, VDJcraft demonstrated superior accuracy in IGH-VDJ reconstruction, achieving a recall of 0.90 and a precision of 0.96 In comparison, TRUST4 showed substantially lower performance, with a recall of 0.581 and a precision of 0.678 while LymAnalyzer achieved a recall of 0.61 and a precision of 0.46. For immunoglobulin light-chain VJ detection, VDJcraft maintained high performance, with a recall of 0.98 and a precision of 0.91. In contrast, TRUST4 achieved a recall of 0.56 and a precision of 0.83, and LymAnalyzer achieved a recall of 0.77 and a precision of 0.75. VDJcraft also outperformed the other tools in T-cell receptor V(D)J detection. For TRA VJ recombination, VDJcraft achieved a recall of 1.00 and a precision of 0.97, while for TRB VDJ recombination, it achived a recall of 0.88 and a precision of 0.90. In comparison, TRUST4 and LymAnalyzer showed reduced accuracy for TCR V(D)J detection, with TRUST4 achieving a maximum recall of 0.68 and a precision of 0.73, and LymAnalyzer showing substantially lower precision, dropping to 0.37 (**Figure 2**).

**Figure 2.**
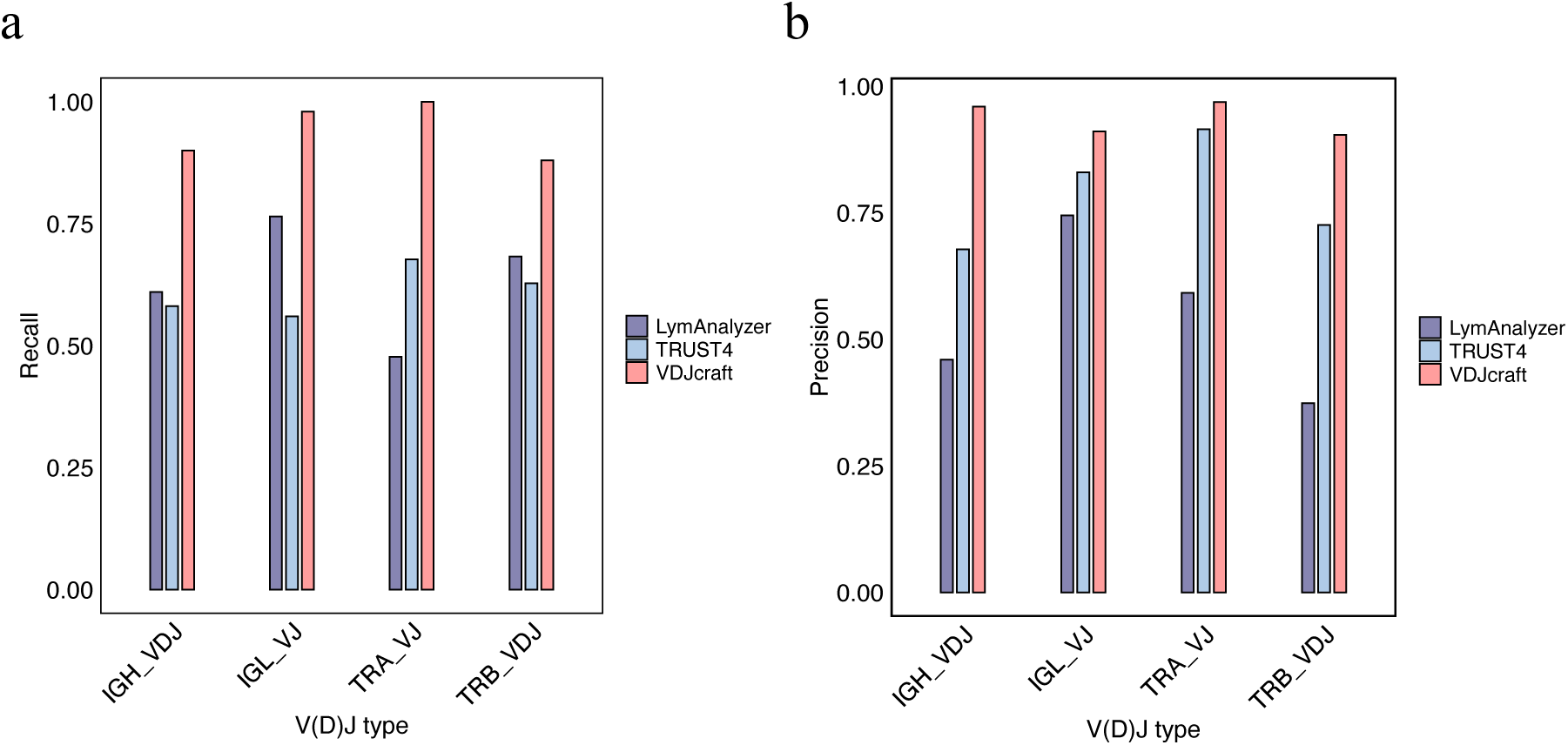
Comparison of V(D)J recombination detection by VDJcraft, TRUST4, and LymAnalyzer on simulated PacBio datasets. Comparison of recall (**a**) and precision (**b**) for V(D)J recombination detection across different immune receptor types using simulated PacBio datasets. Performance of VDJcraft is compared with TRUST4 and LymAnalyzer for IGH VDJ, IGL VJ, TRA VJ, and TRB VDJ recombination.

To further assess detection performance on different types of long reads, we compared two leading long-read sequencing platforms: PacBio and Oxford Nanopore Technologies. Simulated long-read datasets were also generated using PBSIM^34^ and Badread^30^ to model the sequencing characteristics. The performance of VDJcraft was then evaluated separately on simulated datasets from each platform. The results demonstrated that PacBio-based simulations achieved higher recall and precision than Nanopore-based simulations. For TRA VJ detection, PacBio achived an F1 score of 0.98, whereas Nanopore achived an F1 score of 0.93. PacBio also demonstrated higher accuracy in IGH VDJ detection, with a recall of 0.90, a precision of 0.96, and an F1 score of 0.93, compared with a recall of 0.85, a precision of 0.88, and an F1 score of 0.86 for Nanopore (**Table 2**). These findings are consistent with the advantages of PacBio HiFi sequencing, which provides high-fidelity base calling through circular consensus sequencing. Conversely, the comparatively lower performance observed for Nanopore data reflects its higher intrinsic error rates, particularly in homopolymeric regions and in contexts involving base-modification signals, despite its strengths in read length and real-time sequencing capability.

**Table 2.** Performance of VDJcraft on simulated PacBio and Nanopore long-read sequencing datasets.

|  | PacBio |  |  | Nanopore |  |  |
| --- | --- | --- | --- | --- | --- | --- |
|  | Recall | Precision | F1 score | Recall | Precision | F1 score |
| IGH(VDJ) | **0.90** | **0.96** | **0.93** | 0.85 | 0.88 | 0.86 |
| IGK(IGL)VJ | **0.98** | **0.91** | **0.94** | **0.98** | 0.84 | 0.91 |
| TRAVJ | **1.00** | **0.97** | **0.98** | **1.00** | 0.88 | 0.93 |
| TRBVDJ | **0.88** | **0.90** | **0.89** | 0.87 | 0.86 | 0.90 |

Results comparing the accuracy of VDJcraft in V(D)J recombination detection on simulated PacBio and Nanopore sequencing datasets. Recall, precision, and F1 scores are reported for each receptor type, with bold values indicating higher performance.

### Comparison of V(D)J detection between VDJcraft and TRUST4 in 12 HGSVC samples

The Human Genome Structural Variation Consortium (HGSVC) generated PacBio Iso-Seq data to profile full-length transcripts, enabling comprehensive characterization of gene expression and isoform diversity across individuals^26^. To further investigate the capability of VDJcraft to real long-read transcriptomic data, we applied it to 12 PacBio Iso-Seq datasets from the HGSVC and compared the resulting V(D)J profiles with those obtained from matched high-coverage short-read RNA-seq datasets.

TRUST4 was selected as the short-read comparator because of its relatively better performance in simulated benchmarks and its widely adopted in short-read V(D)J recombination analysis. However, inherent challenges associated with short-read sequencing—such as read length, fragmented coverage, ambiguous assembly in repetitive or hypermutated regions, and reduced confidence for low-support events, —necessitated the implementation of a stringent filtering strategy. Specifically, low-confidence calls with insufficient supporting read counts were excluded for comparison to reduce potential false positives.

Comparisons were conducted across three categories to assess the performance and reliability of VDJcraft in detecting V(D)J recombination events: (1) all recombination events identified across the dataset irrespective of quality thresholds, (2) short-read results filtered to exclude low clonotype frequency based on low supporting reads, and (3) putative novel events identified uniquely by VDJcraft. This multi-level approach enabled a comprehensive assessment of detection capabilities across sequencing technologies.

When comparing individual gene segment calls, VDJcraft and TRUST4 demonstrated approximately 60% concordance for V genes, 70% in D genes, and 80% in J genes **(Supplementary Figure 2)**. This observation is consistent with simulation results indicating lower recall of TRUST4 in detecting long V-genes. After applying a threshold of > 0.1% clonotype frequency based on supporting reads in TRUST4, VDJcraft captured up to 100% of the high-confidence V-gene calls detected by TRUST4 and additionally detected novel V gene assignments. Concordance for D- and J-gene calls also increased, reaching approximately 85% and 93%, respectively (**Figure 3a, b**). For VJ recombination detection, initial concordance ranged from 20-60% when low-support TRUST4 calls were included. After filtering to retain only high-confidence short-read events, agreement increased substantially, reaching up to 90%, with most samples clustering near 80% concordance **(Supplementary Figure 2).** The concordance after filtering was significantly increased (Mann-Whitney U test, P = 1.55 × 10⁻⁴) (**Figure 3c**), highlighting the strong influence of read-support quality on V(D)J detection.

**Figure 3.**
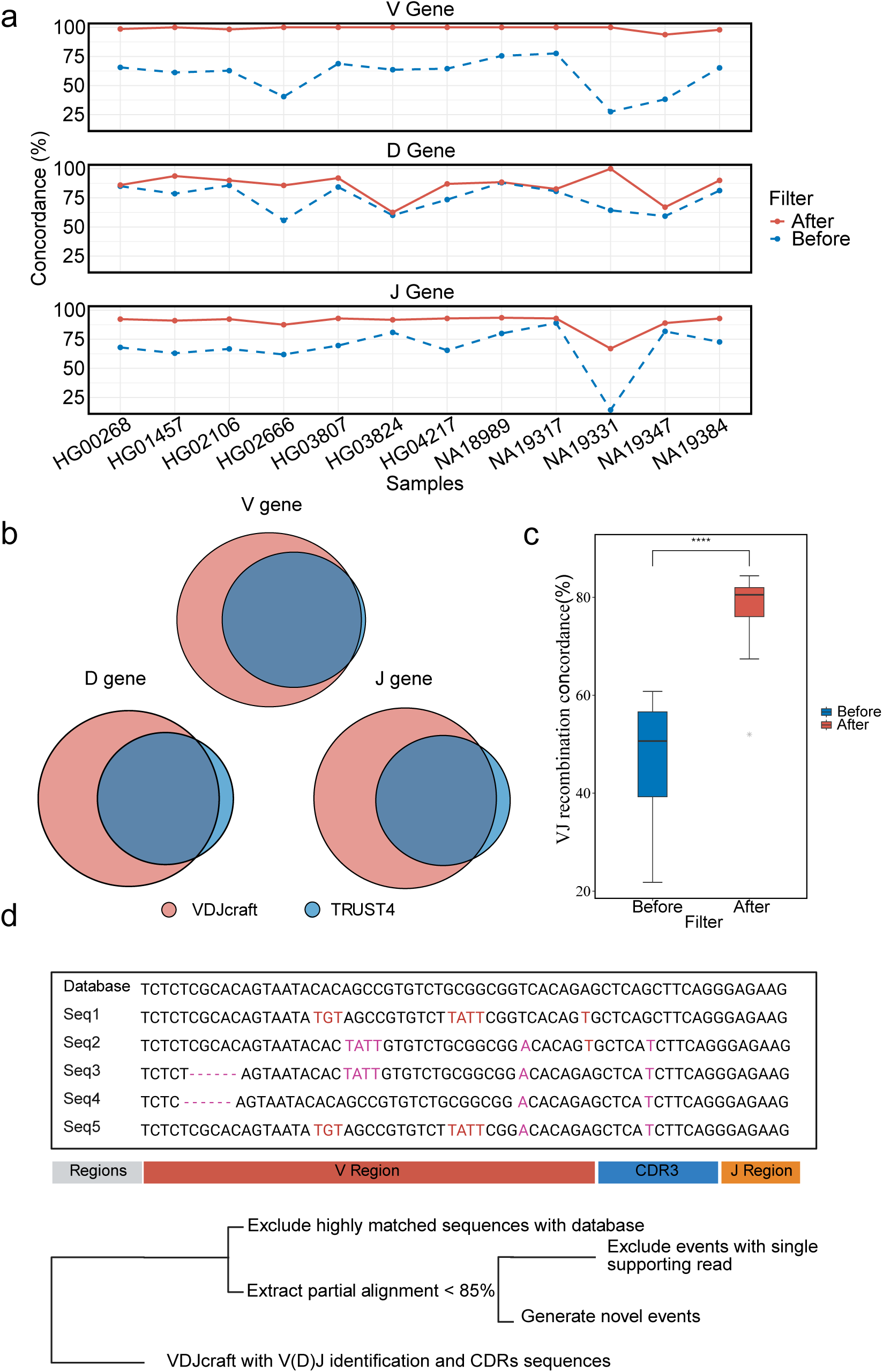
Comparison of VDJcraft and TRUST4 on HGSVC long-read and short-read datasets. a, Comparison of V, D, and J gene-segment calls between VDJcraft (long-read) and TRUST4 (short-read) before and after filtering low-support TRUST4 calls. b, Venn diagrams showing overlap in V, D, and J gene calls between the two methods after filtering out low supporting reads. c, Concordance of VJ recombination calls before and after filtering; concordance increases significantly following removal of low supporting TRUST4 events (Mann-Whiteney U test, P = 1.55 × 10⁻⁴). d, Workflow illustrating the strategy used by VDJcraft to identify putative novel events, including relaxed-identity alignment screening and read-support filtering.

For the detection of complementarity-determining region 3 (CDR3) sequences, VDJcraft employs the same IMGT-based positional framework used by TRUST4, defining the CDR3 region from the conserved motif at the end of the V gene to that at the beginning of the J gene. We compared amino-acid CDR3 sequences derived from long-read (VDJcraft) and short-read (TRUST4) analyses from the 12 HGSVC samples. Detailed comparison of the top 50 TRUST4 CDR3 calls revealed high concordance in light-chain CDR3, observed in heavy-chain CDR3 **(Supplementary Table 2).** These findings highlight that while long-read sequencing can recover nearly all sequences identified by short-read approaches, it additionally captures sequences that short-read reconstruction may miss or misassemble, providing a more comprehensive representation of CDR3 diversity.

In total, 31 putative novel events were identified across the 12 HGSVC samples (**Supplementary Table 3**). To ensure high-quality detection, we focused on reads exhibiting optimal alignment characteristics and selected candidate novel events based on inexact local alignments—specifically, sequences in which < 85% identity to the IMGT database, and at least two supporting reads to minimize false positives (**Figure 3d**). These novel events were predominantly located within the IGHV and IGLV gene regions, a distribution that may reflect the high frequency of somatic hypermutation (SHM) in these loci. In line with previous observations from the TRUST4 algorithm, our pipeline also revealed a higher density of SHMs in the IGH locus compared to the IGK and IGL loci **(Supplementary Figure 3)**. This enrichment supports the central role of the immunoglobulin heavy chain in shaping antigen-binding affinity, highlighting its evolutionary and functional importance in adaptive immune responses^21^. Our observed detection rate (approximately 1.2 x 10^-4^) is consistent with the well-established frequency of somatic hypermutation, which typically ranges from 10⁻³ to 10⁻⁶ mutations per base pair per generation^35,36^. Collectively, these findings demonstrate the capability of VDJcraft to uncover previously unannotated V(D)J recombination events in long-read transcriptomic data and highlight its potential to not only enhance the resolution of immune repertoire analyses but also contribute to the continuous refinement and expansion of the IMGT database by incorporating novel, biologically relevant sequences.

### Cross-validation of VDJcraft and its application to explore immune infiltration in infectious diseases

To comprehensively validate the accuracy and investigate the diverse range of applications of VDJcraft, we applied the tool to analyze datasets generated using the recently developed FLAIRR sequencing technology which uses targeted long reads sequencing to amply immunoglobulin heavy chain transcripts^32^. Specifically, our study employed VDJcraft to detect and characterize IgG subclasses as well as V(D)J recombination events across two distinct sample sets: (1) ten IgG FLAIRR samples derived from healthy individuals, and (2) four longitudinal samples from a single COVID-19 patient, collected at four time points representing different stages of infection and immune response—Day 1, Day 4, Day 8, and Day 13.

For the ten IgG FLAIRR samples, we employed VDJcraft to identify IGHG subclass distributions across seven types of IGHV subclasses. VDJcraft recapitulated the IGHG distribution across IGHV subclasses, showing a trend consistent with the findings reported in the FLAIRR study **(Supplementary Figure 4),** thereby validating its reliability. Furthermore, VDJcraft successfully reproduced the top IGHV gene enrichment in the representative sample (1013) by FLAIRR-seq, with the enriched IGHV subclass distribution aligning precisely with the previously reported data **(Supplementary Figure 5)**. These findings collectively highlight VDJcraft’s robustness in accurately characterizing V(D)J recombination and its potential to enhance our understanding of immunoglobulin diversity.

To demonstrate that VDJcraft can be utilized for disease samples to explore V(D)J recombination pattern that stimulated by specific disease. We applied VDJcraft to analyze samples collected from COVID-19 patients across different recovery days to assess immune system dynamics. Our analysis revealed that IGHV enrichment patterns varied significantly from day 1 to day 13 of recovery. Notably, we identified that specific heavy-chain V genes, such as IGHV1-2_02, IGHV3-30-5_01, IGHV4-39_01, exhibited functional enrichment during recovery day 1 and day 4, then gradually reduced in day 8 and day 13, underscoring their potential importance in the immune response to COVID-19 (**Figure 4a**). Through VDJcraft, we also explored IGH-VDJ recombination patterns and clonotypes over time. A remarkable finding was the significant peak in recombination involving IGHV3-7, IGHD6-19, and IGHJ5-02 on day 4 (**Figure 4b**). This combination represents a potential clonotype critical for the immune system’s ability to defeat COVID-19 virus. This peak suggests day 4 as a pivotal moment for effective immune activation. Additionally, VDJcraft can also characterize class switch recombination (CSR) events. For this dataset, we observed a notable IGHG1-to-IGHG2 switch on day 4, followed by a gradual return to baseline proportions in subsequent days (**Figure 4c**). This trend emphasizes the unique immunological events occurring on around day 4, marking it as a crucial juncture in the recovery process, consistent with previous finding of mean(SD) duration of symptoms 5.4 (3.8) days^37^ and of immune factor serum level enrichment peak at day 5^38^.

**Figure 4.**
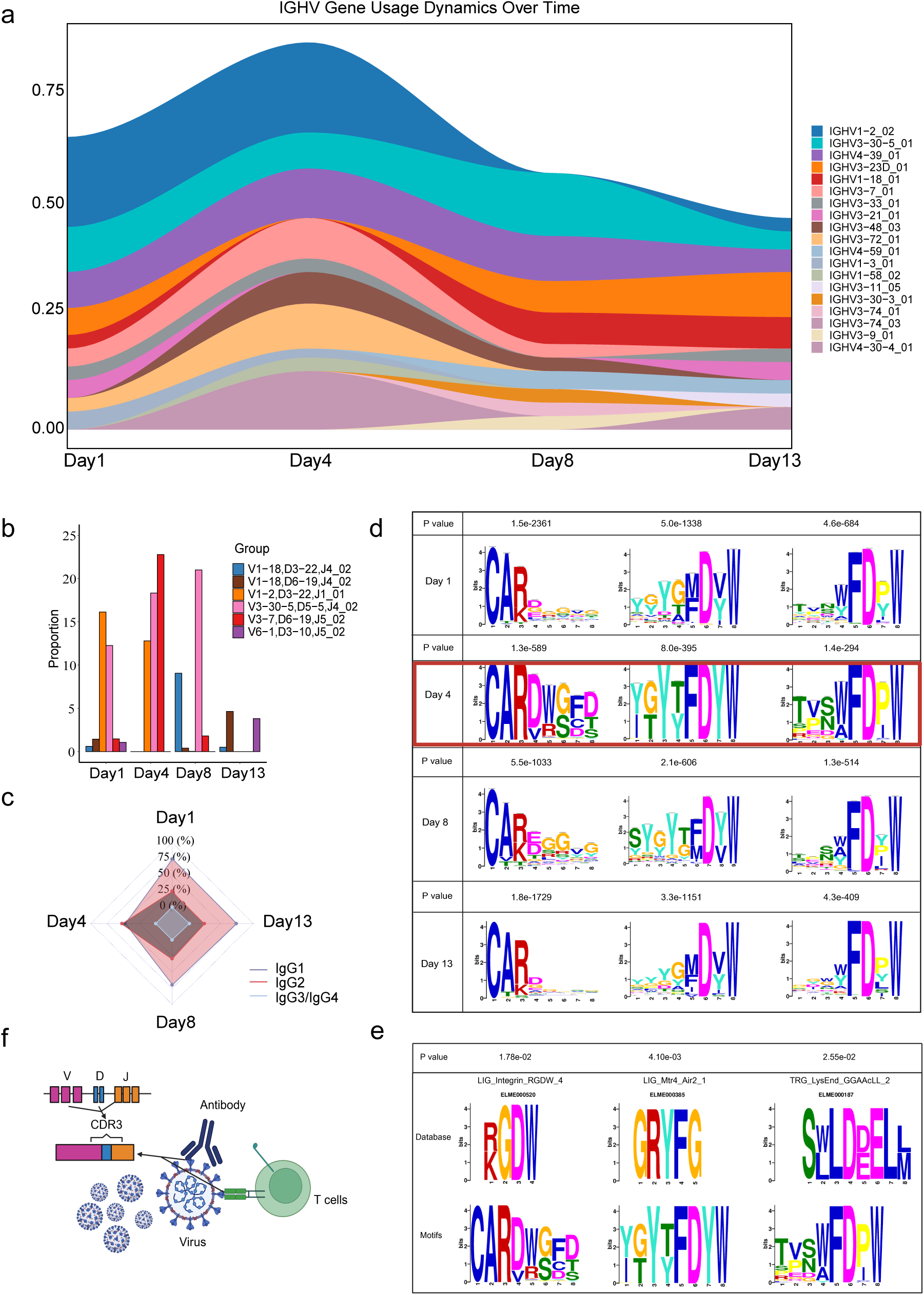
Application of VDJcraft to characterize immune-repertoire dynamics in a longitudinal COVID-19 dataset. a, Ribbon plot showing relative IGHV gene usage on days 1, 4, 8, and 13 after infection; enrichment dynamics during COVID recovery, with IGHV1-2_02 enriched on Day 1 and Day 4, dramatically reduced on Day 8 and Day 13; Day 4 indicates multiple types of IGHV enrichment. b, Dynamics of V(D)J recombination enrichment during COVID recovery, The IGHV1-2–IGHD3-22–IGHJ1_01 combination showed the highest enrichment on Day 1, IGHV3-7–IGHD6-9–IGHJ5_02 peaked on Day 4, and IGHV3-30-5–IGHD5-5–IGHJ4_02 was most enriched on Day 8. By Day 13, enrichment levels were low, suggesting a return of functional immune components to baseline status. c, IgG class switch dynamics during COVID recovery, showing a transient peak in IgG2 levels at Day 4, followed by a return to baseline. d. Antigen binding region motifs dynamics during COVID recovery, showing a transient peak in certain motifs at Day 4 and reduction at Day 8, followed by a return to baseline. e, Functions of top enriched motifs of CDR3 indicate cellular repair, viral RNA elimination, and re-establishment of homeostasis. f. Functional mechanism of antigen binding region for immune response to COVID. The CDR3 region plays a pivotal role in recognizing viral antigens and facilitating their presentation to T cells for clearance.

To explore the potential functional implications of antigen binding domain, We analyzed the amino acid (AA) composition within the CDR3 region using VDJcraft. During the recovery process from COVID-19, the three most significantly enriched amino acid motifs were identified and analyzed. Notably, three dominant motifs enrichment (motif: CARDWGFD; motif: YGYTFDYW; motif: TVSWFDPW) peaked at day 4, gradually reduced on day 8 and returned back to baseline on day13 (**Figure 4d**). These findings suggest the presence of specific amino acid alterations and enrichment dynamics that may play a functional role in immune recovery processes. This observation aligns with broader immunological shifts, particularly those occurring on day 4, and highlights the dynamic nature of the immune repertoire during recovery. To further elucidate the significance of these motifs, we compared the identified motifs against a functional motif database using the MEME Suite comparison module^33^. Remarkably, the motif CARDWGFD demonstrated a significant enrichment to the ‘LIG_Integrin_RGDW_4’ motif in the database^39^, suggesting the presence of an integrin RGD-type binding site^40^. Subunits within this motif can modulate binding specificity and affinity, and integrins themselves play central roles in cell adhesion and communication. Dysregulation of integrin signaling is implicated across a wide range of diseases. Notably, several viruses—including foot-and-mouth disease virus, HIV, West Nile virus, and HPV-16—harbor RGD-like motifs in their proteins, enabling attachment to host-cell integrins and facilitating viral entry^41,42^. Studies have reported that RGD motif SARS-CoV-2 Spike induces TGF-β signaling and downregulates interferon^40^. Motif – TVSWFDPW, corresponds to a Trans-Golgi Network–Endosome–Lysosome sorting signal. The trans-Golgi network is a major hub for membrane protein sorting, particularly near the C-terminus of cargo proteins, where these signals interact with GGA adaptor proteins to recruit clathrin. This motif is located in the hinge region of GGA1 and GGA3 and binds the N-terminal VHS domains of these GGAs^43^. The interaction of such sorting signals with adaptor proteins such as the GGAs (Golgi-localized, gamma ear-containing, ARF-binding protein) is responsible for the targeting of the receptors^44^. Motif – YGYTFDYW, matches the Mtr4–Air2 interaction site, which participates in nuclear RNA surveillance and exosome-mediated degradation of aberrant RNAs^45,46^. These results indicate that recovery from COVID-19 requires both coordinated clearance of the remnants of infection and restoration of normal cellular organization. Membrane-sorting pathways help the cell physically traffic and dispose of viral components, while RNA-degradation pathways eliminate viral RNA and damaged host transcripts. Together, these processes underpin fundamental mechanisms that enable cellular repair, normalization of immune function, and re-establishment of homeostasis during infection recovery (**Figure 4e**).

In summary, VDJcraft provided a comprehensive characterization of the dynamic immunological landscape during the recovery phase of COVID-19. Day 4 emerged as a pivotal time point, marked by pronounced alterations in IGHV-D-J recombination patterns, class-switch recombination events, and the amino acid composition of the CDR3 region. These findings offer critical insights into the adaptive immune response to SARS-CoV-2 infection, enhancing our understanding of the mechanisms underlying immune recovery (**Figure 4f**). Moreover, this study underscores the utility of VDJcraft as a robust tool for dissecting complex immune repertoires across diverse health and disease contexts.

### Runtime and memory usage of VDJcraft

To evaluate the performance of VDJcraft, TRUST4, and LymAnalyzer on long-read sequencing data, we tested each tool using the HGSVC sample HG00268. VDJcraft successfully completed a comprehensive characterization of V(D)J recombination using the long-read dataset within 4 hours and 50 minutes, utilizing a peak memory of 19.769 GB. These results highlight the robustness and efficiency of VDJcraft for accurate V(D)J characterization from long-read sequencing data **(Supplementary Table 1).**

## Discussion

We have developed a long-read transcriptome-based V(D)J recombination detector named VDJcraft. This tool features with comprehensively characterization of V(D)J recombination based on third generation sequencing technology. Based on simulation analysis, real-data comparison of HGSVC samples, VDJcraft demonstrated better performance than short-read bioinformatic tools. Its improved performance is driven by several key design innovations: (1) **Long-read–enabled resolution** — As the first V(D)J analysis tool built specifically for long-read transcriptomic data, VDJcraft fully captures rearranged VDJC regions, providing complete and highly accurate reconstruction. (2) **Dual-pass alignment strategy** — An integrated two-pass alignment approach supports both genome alignment and immune-gene annotation with tunable parameters, ensuring robust detection across diverse sequence contexts. (3) **Targeted D-gene alignment** — A specialized alignment mode enhances the detection of short, highly diverse D genes, improving assignment accuracy even in challenging cases. (4) **CDR3 identification** — As the key determinant of immune receptor diversity and antigen specificity^47,48^. the CDR3 region is precisely captured in VDJcraft using conserved flanking motifs. This enables reliable extraction of full CDR3 sequences and provides rich information for downstream analyses of antigen-binding features. (5) **Built-in error correction** — VDJcraft enhances result reliability by correcting potential sequencing errors, particularly in regions supported by only one or two reads. This increases confidence in low-coverage reconstructions and improves robustness for downstream immunogenomic interpretation. (6) **Broad applicability and efficient performance** — VDJcraft includes multiple functional modules supporting both PacBio and ONT data. It’s also suitable for studies in both healthy and disease contexts. Additionally, the tool is optimized for computational efficiency, demonstrating reduced runtime and memory usage across diverse analytical workflows.

Previous literatures have reported that using next-generation sequencing to effectively capture V(D)J recombination can reflect the immune response and infiltration in infection^49^ as well as tumor environment^3,50^. V(D)J recombination, as a critical biological process, underpins the development and diversity of the adaptive immune system. Understanding this process is essential for elucidating disease mechanisms, immune responses, and potential therapeutic targets. Somatic hypermutation (SHM), which introduces random mutations during B-cell maturation to increase the affinity and specificity of antigen recognition, represents a crucial process for comprehensive characterization^51^. With the advent of third-generation sequencing technologies, it has become possible to unravel the intricate patterns of V(D)J recombination at an unprecedented resolution, offering valuable insights into its role in health and disease status. This study demonstrates the efficacy of VDJcraft, a first novel computational tool based on third generation sequencing, in identifying V(D)J recombination patterns and immune infiltration that contribute to immune responses, validated in the application context of COVID-19. Previous research has shown that B-cell receptors (BCRs) in COVID-19 patients tend to utilize different V gene segments compared to healthy controls. Notably, the complementarity-determining region 3 (CDR3) sequences of heavy chains in clonal BCRs from patients exhibit greater convergence relative to those from healthy individuals^49,52^. Additionally, these studies have reported an increase in IgG and IgA isotypes in COVID-19 patients, indicative of an adaptive immune response ^53^. Our analysis corroborates these findings, highlighting a significant enrichment of the IGHV3-30 and IGHV1 gene families in COVID-19 patients^54^. Moreover, de novo computational designs of high-affinity antibody variable regions (Fv) have shown promise in targeting the most solvent-exposed ACE2-binding residues of the SARS-CoV-2 spike receptor-binding domain (RBD) protein. By recombining VDJ genes and introducing mutations, researchers generated high-affinity variable regions targeting spike protein epitopes, underscoring the diversity and plasticity of V(D)J recombination in generating effective immune responses against viral pathogens^55^. Dysfunctional V(D)J recombination has been implicated in various diseases, including autoimmune disorders and immunodeficiencies. For instance, monoallelic mutations in LIG4, a pivotal enzyme responsible for sealing DNA breaks during V(D)J recombination, can lead to autoimmunity and immunodeficiency through haploinsufficiency^56^. Errors at any step of the V(D)J recombination process can result in abnormalities^57,58^.Additionally, studies have reported that IGHV1-69, a germline gene segment, is prominently utilized in antiviral monoclonal antibodies (mAbs) against pathogens such as influenza virus^59^, HIV^60^, SARS-CoV^61^, Hendra virus^62^, dengue virus^63^, and Nipah virus^62^. This highlights the conserved role of specific germline gene segments in mounting immune responses to diverse viral infections, underscoring the importance and the urgency of comprehensively and accurately characterizing this process within the immune system.

In addition, third-generation sequencing (TGS) has transformed cancer research by enabling comprehensive analysis of complex genomic events. Its ability to generate long reads that span large, repetitive regions allows precise resolution of structural rearrangements that underlie disease initiation and progression^64^. Recent studies have identified a distinct B-cell subset with a unique class-switch signature and strong effector capacity, underscoring its role in antibody-dependent cytotoxicity and protective immunity such as anti-tumor immunity^3^. Higher T-cell receptor V(D)J read recovery—reflecting greater T-cell infiltration into primary tumors—correlates with poorer overall and disease-free survival and more advanced disease stage^65^. V(D)J recombination generates diverse antigen receptors through tightly regulated DNA rearrangement and repair; disruptions to this process can produce faulty receptors, impair immunity, and increase susceptibility to immunodeficiencies and malignancies^66–69^. These findings collectively underscore the critical importance of comprehensively characterizing V(D)J recombination. Leveraging third-generation sequencing technologies provides unprecedented resolution of B cell and T cell receptor repertoires, enabling deeper insight into adaptive immune diversity, clonal dynamics, and functional responses. Such high-resolution profiling clarifies immune function and dysregulation, reveals mechanisms of disease, and guides precision immunomodulatory strategies for improved prevention and therapy.

Even though VDJcraft shows higher performance and comprehensiveness to identify and analyze V(D)J recombination based on long reads. There are some limitations that currently haven’t been solved or needs to be further improved. First, long reads have higher accuracy but it’s not perfect. Errors caused by sequence platform can be corrected in some level but cannot be fully addressed. Second, this tool is alignment-based, meaning that identification relies on the reference. It may potentially miss novel events that cannot be aligned to the reference, and its performance is inherently limited by the completeness of the reference.

Collectively, VDJcraft is an advanced bioinformatics tool specifically designed for the comprehensive characterization of V(D)J recombination using long-read transcriptome sequencing data. This tool excels in accurately identifying and annotating V, D, J, and C genes that constitute unique clones, and characterizing key sequences such as CDR1, CDR2, and CDR3. By leveraging the advantages of third-generation sequencing technology, VDJcraft addresses several limitations associated with next-generation sequencing based tools, such as incomplete gene detection and reduced sequence resolution in complex immune loci. Extensive benchmarking demonstrates that VDJcraft outperforms short-read-based methods in both simulated and real datasets, providing superior accuracy in detecting V(D)J genes and capturing full-length sequence information. The tool integrates a built-in error correction step, which enhances the reliability of its outputs, particularly in datasets prone to sequencing errors or artifacts. This feature ensures a higher level of resolution in characterizing immune repertoires. A key strength of VDJcraft lies in its computational efficiency. Most of its algorithms are designed with linear computational complexity, which significantly reduces the time and resources required to execute the pipeline. This optimization makes VDJcraft a powerful and practical solution for large-scale studies, enabling researchers to analyze immune repertoires with greater detail and accuracy while maintaining efficiency.

## Acknowledgements

We thank the Human Genome Structural Variation Consortium (HGSVC) for generating and making available the PacBio Iso-Seq and high-coverage Illumina short reads datasets used in this study. We also thank Ryan Melvin, Gerald McGwin, and Hui-Chen Hsu for their helpful comments on the manuscript. Funding was provided by MIRA Award R35GM138212 grant.

## Author information

First author contributes primarily: Kaili Hu

## Contributions

Study concept, planning, and design (KH, ZC), tool development (KH), figure/table generation and summary (KH), tool test (YS, CF, ZP), figure/table refinement (AR, YS, CF, ZP, MG, ZC), drafting of initial manuscript (KH), editing of manuscript (ZC), critical review (all authors) Sources of Support: R35GM138212 (ZC)

