## Supplementary material for "Comprehensive characterization of V(D)J recombination from long-read transcriptomic data with VDJcraft": VDJ_Supplementary_materials_v1_2026.pdf

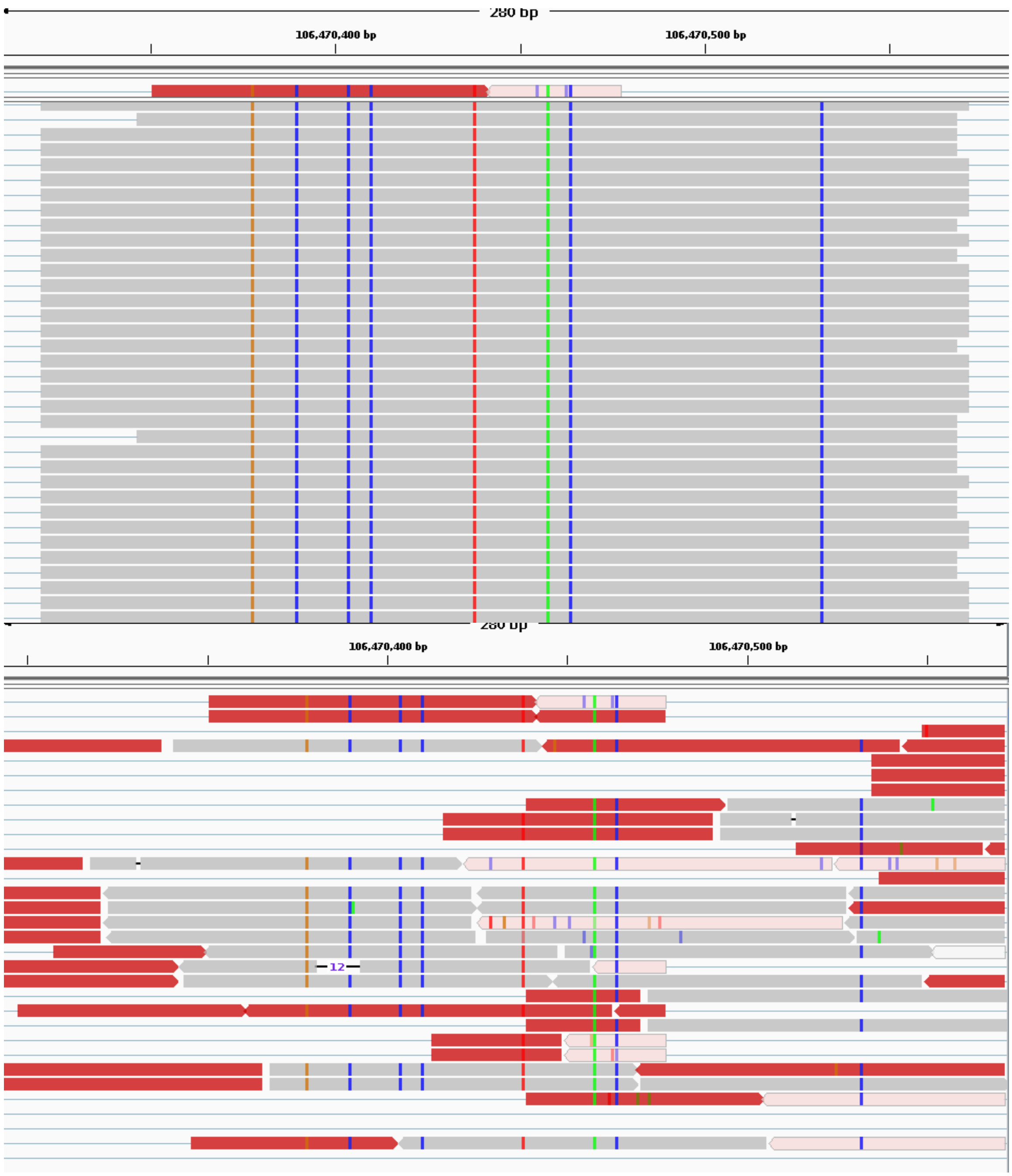


**Supplementary Figure 1. Comparison of IGV view for IGHV3-43 gene between long reads and short reads.** Upper panel shows long reads alignment in IGHV3-43 gene region, below panel shows short reads alignment in IGHV3-43 gene region.

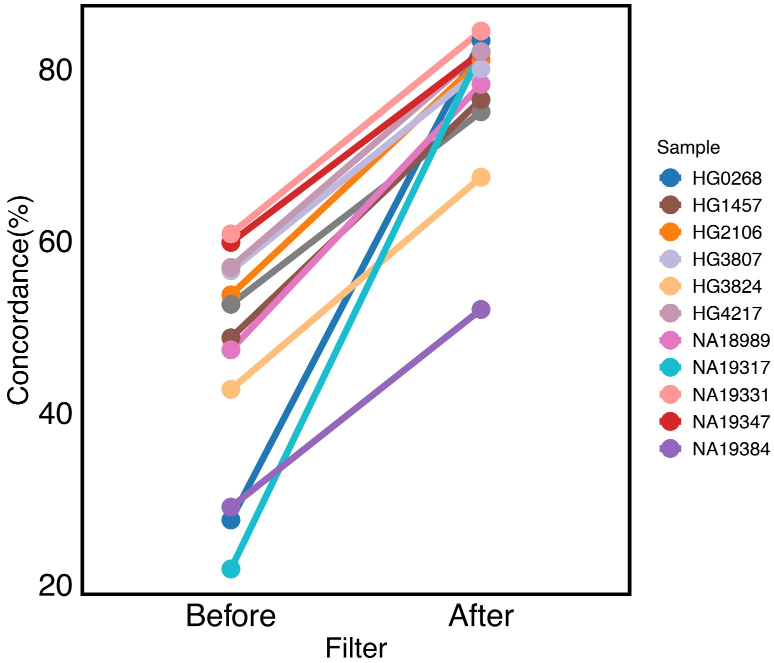


**Supplementary Figure 2. Concordance of VJ recombination between long reads and short reads on HGSVC Samples VDJcraft and TRUST4.** VJ recombination concordance between VDJcraft and TRUST4 increased markedly after filtering out low-supporting clonotypes compared to before filtering.

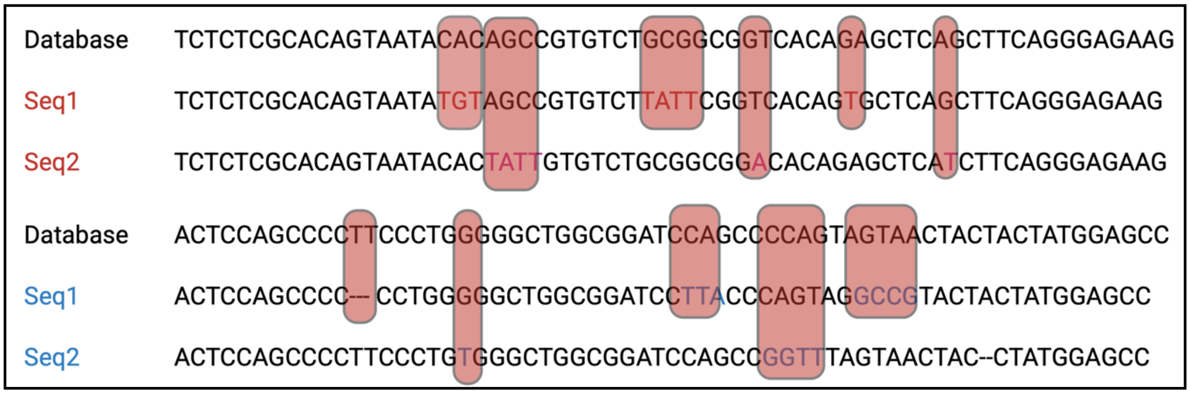


**Supplementary Figure 3. Novel events identified by VDJcraft potentially induced by somatic hypermutation.** Mismatched sequences are highlighted within the sequences, indicating regions where somatic hypermutation has occurred.

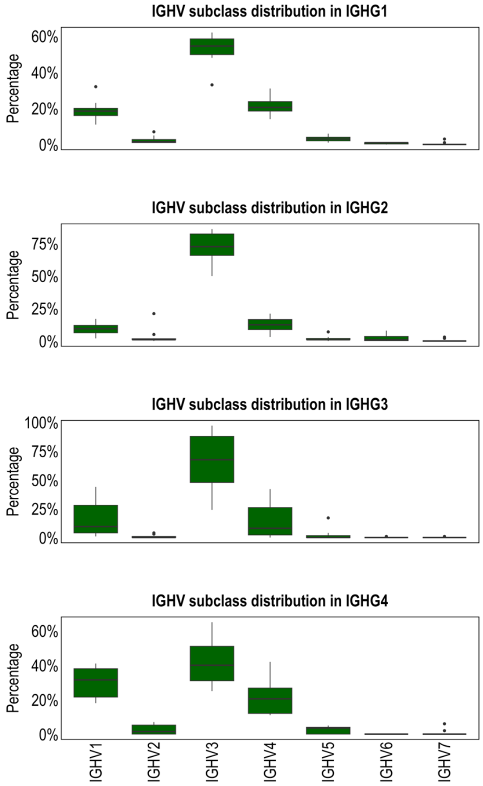


**Supplementary Figure 4. IGHV subclass distribution of 10 FLAIRR samples using VDJcraft consistent with the results shown on previous literature.** The trend of IGHV family usage across different IGHG subclasses detected by VDJcraft is consistent with previously reported findings[1].

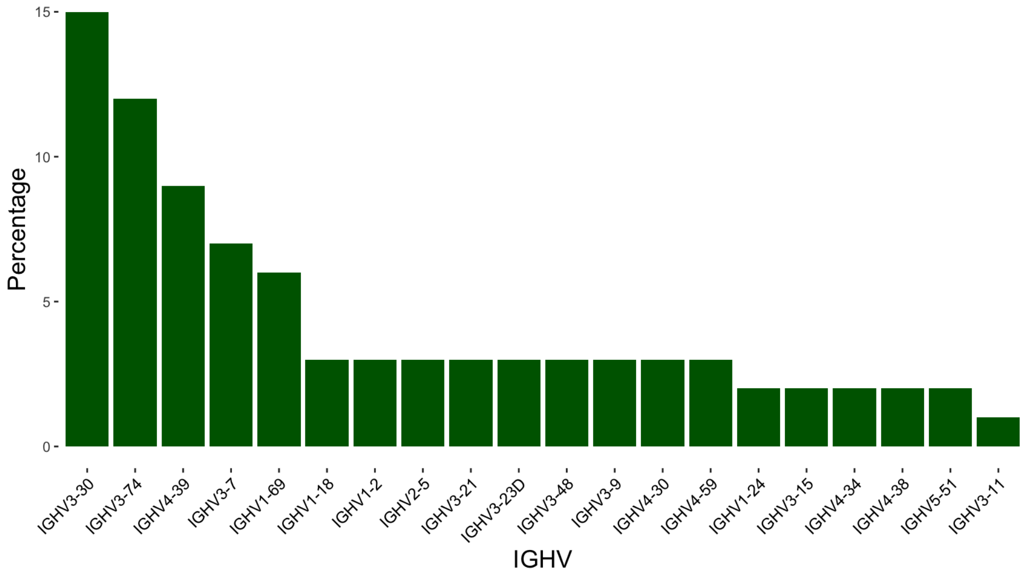


**Supplementary Figure 5. Top 20 IGHV gene enrichment in FLAIRR1013 sample by percentage using VDJcraft consistent with the results shown on previous literature.** IGHV3-30 exhibited the highest usage among IGHV genes in this sample, consistent with previously reported findings[1].

**Supplementary Table 1.** Comparison of VDJ detection on HGSVC sample (HG00268) by VDJcraft. TRUST4, and Lymanalyzer.

| Measurement | VDJcraft | TRUST4 | Lymanalyzer |
|---|---|---|---|
| **Outcome** | Success | Failed | Failed |
| **Time consuming** | 4h 50min | - | 50 hour of runtime or more |
| **Memory usage** | 19.769G | (20 GB memory per CPU; 20 GB CPU per task) | 8 GB memory per CPU; 5 GB CPU per task |
| **Error** | - | Out of memory | - |
| **Output** | Complete report | No output | No output |

**Supplementary Table 2.** Comparison of top enriched CDR3 sequences identified by VDJcraft and TRUST4.

| VDJ_clonotype | VDJcraft_CDR3nt | VDJcraft_CDR3aa | TRUST4_CDR3nt | TRUST4_CDR3aa |
|---|---|---|---|---|
| IGHV7-4-1_02 . IGHJ4_02 | TGTGCGAGAGATGACCCTTACGAGCTTTTGACTGGTAATGACCTCTTT | CARDDPYELLTGNDLF | TGTGCGAGAGATGACCCTTACGAGCTTTTGACTGGTAATGACCTCTTTGACAAGTGG | CARDDPYELLTGNDLFDKW |
| IGLV1-47_02 . IGLJ3_02 | GCAACATGGGATGACAGCCTGAGTGCTTGG | ATWDDSLSAW | TGTGCAACATGGGATGACAGCCTGAGTGCTTGGGTGTTC | CATWDDSLSAWVF |
| IGHV3-23_01 . IGHJ4_02 | TGTGCGAAATGTTCCTGTAAAGCAGGGGCTGACCAACTTGACCACTGG | CAKCSCKAGADQLDHW | TGTGCGAAATGTTCCTGTAAAGCAGGGGCTGACCAACTTGACCACTGG | CAKCSCKAGADQLDHW |
| IGHV4-4_07 . IGHJ3_02 | TGTGCGAGAGTCGGGGATAGTAGTGGTTATTACCCTGATGCTTT | CARVGDSSGYYPDAF | TGTGCGAGAGTCGGGGATAGTAGTGGTTATTACCCTGATGCTTTTGATATCTGG | CARVGDSSGYYPDAFDIW |
| IGKV1-39_01 . IGKJ1_01 | TGTCAACAGAGTTACAGTACCCCTTGG | CQQSYSTPW | TGTCAACAGAGTTACAGTACCCCTTGGACGTTC | CQQSYSTPWTF |
| IGHV1-18_01 . IGHJ4_02 | TGTGCGAGGGGAAAATATAGTAGTGGTTGGCCCGCTGACTACTGG | CARGKYSSGWPADYW | TGTGCGAGAACGGCGGTCTACTTCGGGGACTTTGACTCCTGG | CARGTDTVVVTAPIDYW |
| IGLV1-47_02 . IGLJ3_02 | GCAACATGGGATGACAGCCTGAGTGCTTGG | ATWDDSLSAW | TGTGCAGCATGGGATGACAGCCTGAGTGGTCCAGGATGGGTGTTC | CAAWDDSLSGPGWVF |
| IGKV3-15_01 . IGKJ2_01 | TGTCAGCAGTATAATAACTGGCCTCCGTACACTTT | CQQYNNWPPYTF | TGTCAGCAGTATAATAACTGGCCTCCGTACACTTTT | CQQYNNWPPYTF |
| IGHV4-4_07 . IGHJ4_02 | TGTGCGAGGGGGGAGTATTACTATGATAGTAATGGGTATGACTACTGG | CARGEYYYDSNGYDYW | TGTGCGAGAACGGCGGTCTACTTCGGGGACTTTGACTCCTGG | CARTAVYFGDFDSW |
| IGLV2-14_01. IGLJ1_01 | TGCAGCTCATATACAAGCAGCAGCACTTATGTCTTC | CSSYTSSSTYVF | TGCAGCTCATATACAAGCAGCAGCACTCGAGTCTTC | CSSYTSSSTRVF |
| IGHV3-11_01. IGHJ6_02 | TGTGCGAGGACCCCGACTGGAAACAATTATGGTTCAGACTACTACTACTACGGTATGG | CARTPTGNNYGSDYYYYGMW | TGTGCGAGGACCCCGACTGGAAACAATTATGGTTCAGACTACTACTACTACGGTATGGACGTCTGG | CARTPTGNNYGSDYYYYGMDVW |
| IGHV4-34_10. IGHJ4_02 | TGTGCGAGGGGAACAGACACTGTGGTGGTGACTGCTCCTATTGACTACTGG | CARGTDTVVVTAPIDYW | TGTGCGAGAGTGTTACGTCGAGTTACTACGATAGTAGTGGTTTTTGACTACTGG | CARVLRRVTTIVVVFDYW |
| IGHV3-64_07 . IGHJ3_02 | TGCGAGAGCCCCCGCAGCCCTGAGGGCTGCTTT | CESPRSPEGCF | TGTGCGAGAGCCCCCGCAGCCCTGAGGGCTGCTTTTGATATCTGG | CARAPAALRAAFDIW |
| IGKV3-15_01. IGKJ3_01 | TGTCAGCAGTATAATAACTGGCCTTTC | CQQYNNWPF | TGTCAGCAGTATAATAACTGGCCTTTCACTTTC | CQQYNNWPFTF |
| IGLV2-23_01 . IGLJ3_02 | TGCTGCTTATATGCAGGTAACACTGATTGG | CCLYAGNTDW | TGCTGCTTATATGCAGGTAACACTGATTGGGTGTTC | CCLYAGNTDWVF |
| IGKV3-20_01. IGKJ2_01 | TGTCAGCAGTATGGTGGCTCACCCCCGGGGTACACTTT | CQQYGGSPPGYTF | TGTCAGCAGTATGGTGGCTCACCCCCGGGGTACACTTTT | CQQYGGSPPGYTF |
| IGKV3-20_01. IGKJ4_01 | TGTCAGCAGTATTCTGGCTCACAGAGGACTTT | CQQYSGSQRTF | TGTCAGCAGTATTCTGGCTCACAGAGGACTTTC | CQQYSGSQRTF |
| IGHV3-66 . IGHJ6_02 | TGTGCGAGATTCGGCTATAGCAGCCCACCCAGACCGTTCTACTACTACTACGGTATGG | CARFGYSSPPRPFYYYGMW | TGTGCGAGATTCGGCTATAGCAGCCCACCCAGACCGTTCTACTACTACGGTATGGACGTCTGG | CARFGYSSPPRPFYYYGMDVW |
| IGHV3-48_01 . IGHJ4_02 | TGTGCGAGCCCAAAGTCAGAGGGATCCAGGGATTACTTT | CASPKSEGSRDYF | TGTGCGAGCCCAAAGTCAGAGGGATCCAGGGATTACTTTGACTACTGG | CASPKSEGSRDYFDYW |
| IGLV1-44_01 . IGLJ1_01 | TGTGCAGCGTGGGATGACAGCCTGAATGGCAATGTCTTC | CAAWDDSLNGNVF | TGTGCAGCGTGGGATGACAGCCTGAATGGCAATGTCTTC | CAAWDDSLNGNVF |
| IGKV3-20_01 . IGKJ1_01 | TGTCAGCAGTATTCTGGCTCACAGAGGACTTT | CQQYSGSQRTF | TGTCAGCAGTATAGTAGCGCACCGTGGACGTTC | CQQYSGSQRTF |
| IGHV3-9_01 . IGHJ3_02 | TGTGTGAAGGTAATACACAGTGCCATTGG | CVKVIHSAIW | TGTGTGAAGGTAATACACAGTGCCATTGGTGCTTTTGATATCTGG | CVKVIHSAIGAFDIW |
| IGLV2-14_01 . IGLJ3_02 | TGCAGCTCATATACAAGCAGCAGCACTTATGTCTTC | CSSYTSSSSWVF | TGCAGCTCATATACAAGCAGCAGCTCTTGGGTGTTC | CSSYTSSSSWVF |
| IGKV3-15_01 . IGKJ4_01 | TGTCAGCAGTATAATAAATGGCCGCTCACTTT | CQQYNKWPLTF | TGTCAGCAGTATAATAAATGGCCGCTCACTTTC | CQQYNKWPLTF |
| IGHV3-7_01 . IGHJ5_02 | TGTGCGGGACTCAGCTACATGGCATTT | CAGLSYMAF | TGTGCGGGACTCAGCTACATGGCATTTGACCTCTGG | CAGLSYMAFDLW |
| IGLV2-23_02 . IGLJ3_02_IGLV 2-11_01 IGLJ3_02) | TGCTGCTCATATGCAGGCAGCTACACTTTCGGGGTGTTC | CCSYAGSYTFGVF | TGCTGCTCATATGCACGTAGTAGCACCTGGGTGTTC | CCSYARSSTWVF |
| IGHV3-15_01 . IGHJ4_02 | TGTACCACAGGGTATGGCCCTGACTACTGG | CTTGYGPDYW | TGTAGTACAGATGATTACGATATTTTGACTGGTTATTCCGATTACTGG | CSTDDYDILTGYSDYW |
| IGLV1-44_01 . IGLJ3_02 | TGTGCAGCGTGGGATGACAGCCTGAATGGCAATGTCTTC | CAAWDDSLNGNVF | TGTGCAGCATTGGATGACAGCCTGAATGGCTGGGTGTTC | CAALDDSLNGWVF |
| IGLV3-21_03 . IGLJ3_02 | TGTCAGGTGTGGGATAGTAGTTGGGGCGTCTTC | CQVWDSSWGVF | TGTCAGGTGTGGGATGGAGGTGTTGCCTGGGTGTTC | CQVWDGGVAWVF |
| IGHV1-2_02 . IGHJ5_02 | TGTACGACATATCAGGACAACCAGTGGTTGCCCCCGTTC | CTTYQDNQWLPPF | TGTACGACATATCAGGACAACCAGTGGTTGCCCCCGTTCGACCCCTGG | CTTYQDNQWLPPFDPW |
| IGLV6-57_04. IGLJ3_02 | TGTCAGTCTTATGATACCAGCAATCTCGTGTTC | CQSYDTSNLVF | TGTCAGTCTTATGATACCAGCAATCTCGTGTTC | CQSYDTSNLVF |
| IGKV1-5_01. IGKJ1_01 | TGCCAACAGTATAATAGTTATTGG | CQQYNSYW | TGCCAACAGTATAATAGTTATTGGACGTTC | CQQYNSYWTF |
| IGHV3-9_01 . IGHJ6_02 | TGTGTTAGAGCAGTGCCCGGAGAGGGCGGTATGG | CVRAVPGEGGMW | TGTGTTAGAGCAGTGCCCGGAGAGGGCGGTATGGACGTCTGG | CVRAVPGEGGMDVW |
| IGHV4-4_07 . IGHJ4_02 | TGTGCGAGGGGGGAGTATTACTATGATAGTAATGGGTATGACTACTGG | CARGEYYYDSNGYDYW | TGTGCGAGGAGTCTTGATTGGCAGTATCCCTTTGACTTCTGG | CARSLDWQYPFDFW |
| IGLV1-44_01 . IGLJ3_02 | TGTGCAGCATTGGATGACAGCCTGAATGGCTGG | CAALDDSLNGW | TGTGCAGCATGGGATGACAGTTTGAAGGGTTGGGTGTTC | CAAWDDSLKGWVF |

| | | | | |
|---|---|---|---|---|
| IGHV1-2_02 . IGHJ5_02 | TGTACGACATATCAGGACAACCAGTGGTTGCCCCCGTTC | CTTYQDNQWLPPF | TGTACGACATATCAGGACAACCAGTGGTTGCCCCCGTTCGACCCCTGG | CTTYQDNQWLPPFDPW |
| IGLV6-57_04 . IGLJ3_02 | TGTCAGTCTTATGATAGCAGCGGCTGG | CQSYDSSGW | TGTCAGTCTTATGATACCAGCAATCTCGTGTTC | CQSYDTSNLVF |
| IGKV1-5_01 . IGKJ1_01 | TGCCAACAGTATAATAGTTATTGG | CQQYNSYW | TGCCAACAGTATAATAGTTATTGGACGTTC | CQQYNSYWTF |
| IGHV3-9_01 . IGHJ6_02 | TGTGTTAGAGCAGTGCCCGGAGAGGGCGGTATGG | CVRAVPGEGGMW | TGTGTTAGAGCAGTGCCCGGAGAGGGCGGTATGGACGTCTGG | CVRAVPGEGGMDVW |
| IGHV4-4_07 . IGHJ4_02 | TGTGCGAGGGGGGAGTATTACTATGATAGTAATGGGTATGACTACTGG | CARGEYYYDSNGYDYW | TGTGCGAGGAGTCTTGATTGGCAGTATCCCTTTGACTTCTGG | CARSLDWQYPFDFW |
| IGLV1-44_01 . IGLJ3_02 | TGTGCAGCATTGGATGACAGCCTGAATGGCTGG | CAALDDSLNGW | TGTGCAGCATGGGATGACAGTTTGAAGGGTTGGGTGTTC | CAAWDDSLKGWVF |
| IGKV3-15_01. IGKJ2_01 | TGTCAGCAGTATAATAACTGGCCTCCGTACACTTT | CQQYNNWPPYTF | TGTCAGCAGTATAATAACTGGCCGTACACTTTT | CQQYNNWPYTF |
| IGHV3-7_01 . IGHJ4_02 | TGTGCGAGAGATCAGTGGCGGATCTTT | CARDQWRIF | TGTGCGAGAGATCAGTGGCGGATCTTTGACTACTGG | CARDQWRIFDYW |
| IGLV3-9_01 . IGLJ1_01 | TGTCAGGTGTGGGACGCCGGCACTGCAGGGTATGTCTTC | CQVWDAGTAGYVF | TGTCAGGTGTGGGACGCCGGCACTGCAGGGTATGTCTTC | CQVWDAGTAGYVF |
| IGKV1-33_01 . IGKJ5_01 | TGTCAACAGTATGATAATCTCCCCCCGATCACCTTC | CQQYDNLPPITF | TGTCAACAGTATGATAATCTCCCCCCGATCACCTTC | CQQYDNLPPITF |
| IGHV2-5_02 . IGHJ4_02 | TGTGCACACTTTGGCCAAGTCTACTTT | CAHFGQVYF | TGTGCACACTTTGGCCAAGTCTACTTTGACTACTGG | CAHFGQVYFDYW |
| IGHV3-66_01 . IGHJ6_02 | TGTGCGAGAGATGGGTGGTTAGGGGCCGGGGGTTT | CARDGWLGAGGF | TGTGCGAGAGATGGGTGGTTAGGGGCCGGGGGTTTGGACGTCTGG | CARDGWLGAGGLDVW |
| IGHV4-61_12 . IGHJ5_02 | TGTGCGAGACATTTACAGATTTTTGCCTGG | CARHLQIFAW | TGTGCGAGACATTTACAGATTTTTGCCTGGTTCGACCCCTGG | CARHLQIFAWFDPW |
| IGHV1-18_01. IGHJ4_02 | TGTGCGAGAGGGAGAGACCCAGCTGCCGATCTTGACTTC | CARGRDPAADLDF | TGTGCGAGAGGGAGAGACCCAGCTGCCGATCTTGACTTCTGG | CARGRDPAADLDFW |
| IGLV6-57_04 . IGLJ3_02_IGHV3-23D_01 IGHJ1_01) | TGTGCGAACCAACAGTCCCAAAAACTACGGGAATACTTC | CANQQSQKLREYF | TGTGCGAACCAACAGTCCCAAAAACTACGGGAATACTTCCAGCACTGG | CANQQSQKLREYFQHW |

**Supplementary Table 3.** Novel genes identified by VDJcraft potentially enrich IMGT database.

| count | Gene_class | Novel_seq |
|---|---|---|
| 8 | IGHV1-18* | CTCTCGCACACTAATATAAGGCCGTGTCGTCAGATCTCAGGCTCCTCAGGTCCATGTAGAGTGTAGTCGCGGACGTGTCTCTGGTCAAGGTGACTCTGCCCTAGAACTGCTGTGTGTAGGTTGTGCTGCCA |
| 11 | IGHV1-18* | TGGCAGCACAACCTACACACAGCAGTTCTAGGGCAGAGTCACCTTGACCAGAGACACGTCCGCGACTACACTCTACATGGACCTGAGGAGCCTGAGATCTGACGACACGGCCTTATATTAGTGTGCGAGAG |
| 4 | IGHV1-69* | TCTCCCGCACAGTAATATACGGCCGTGTCTTCAGATGTCAGATTCGTTAGTTCCATGTAGACCATGTTTGTCTTCTTTTCCGCGGTCATTGTGACTCTGCCGTGGAATTTCTGTGCGTAGTGTGGAATATTAA<br>GTAGAGGGATGATCCTTCCCATCCACTCAAGTCCATGTCCTGGGGCCTGTCGCACCCAGCTGATACTATTACTGATGAGGGTGCCGCCAGAAGCCTTACAG |
| 2 | IGHV1-8* | CAGGTGCAGGTGTTGCATTCTGGGGCTGAGGTGAAGAAGCCTGGGGCCTCAGTGCAGGTCTCCTGGAAGGCTTCTGGATACACCTCCGTCACTTATGGTATCAACTGGGTGCGACAGGCCACTGGACACGG<br>GCTGCGACAGGCCACTGGACAAGGGCTTGAATGCATGGGATGGATGAACCCTAACAGTGGTGGCACAGGCTATGCACAGATTTTCCAGGGCAGAGTCACCATG |
| 2 | IGHV1-8* | TCTCGCACAATAATACGCGGCCGTGTCTTCAAATCTCAGGCTGCTCATCGCCATGCAGCCTGTGTTTATGGAGGTGTTCCTGGCCATGGTGACTCTGCCCTGGAAAATCTGTGCATAGCCTGTGCCACCACT<br>GTTAGGGTTCATCCATCCCATGCATTCAAGCCCTTGTCCAGTGGCCTGTCGCAGCCCGTGTCCAGTGGCCTGTCGCACCCAGTTGATACCATAAGTGACGGA |
| 2 | IGHV3-15* | GTGGTTCAAAAAGACACGCCTGTGTCCTCGGTTTTCAGGCTGTTCATTTCCAAACACAGCGGAGTTTTCGGTTAATCTCTTCAGACGGTGAATCAGGCTTTCACGGGCACAACGTAGTCCGGTGCCCCAACAT<br>CACCTTTTGGTTTTAAGACGGCCAACCCACTCCAGCCCCCTCCCTGCAGCCTGGCGGACCCACCTCATCC--------------GTGAATCCAGAGTCTAC |
| 2 | IGHV3-23D_* | GAGGTACAATTGTTGGATTCTGGGGGAGGCTTGGTACAGCCCTGGGGGGTCCCTGAGACTCTCCTGTTCAGCCGCTGGATTCACGTTTACCAACTTTGCCATGACCTGGGTC-GCCAGGCTCC-GGGAGGGGG<br>CTCGAGTGGGTCTCAACTTTATTCCTAGTGACGATAGAGTAAACTACGATAGAACATACTACACAGACTCCGTGAAGGGCCGGTTCACCATCTCCAGAGAC |
| 2 | IGHV3-33_* | CTCTTGCACATTAATACACAGCCGTGTCCCCGGCTCTCAGGCTGTTCATTTGAAGATACAAGGAGTTCTTGGCATTTTCTCTGGAGATGGTGAATCGGCCCTTCACGGAGCCTGGATCCTCTATGTCACCAG<br>CA-CTT-CCA-ATAGTTGAGACCCATTCCAGACCTTTTCCTGTCGCTTGGCGGACCCAGTGCATGTCGTAGCTACTGAAGGTGAATCCAGAGGCTGCACAGG |
| 8 | IGHV3-48_* | GACGTGCAGCTGGTGGAGTCGGGGGGACACTTGGTCCAGCCgggggggTCCCTGAGGCTCTCCTGTGCAGCCTCTGGATTCAGTTTTAGTAGTTTTTGGATGAGTTGGGTCCGCCAGGCTCCAGGGAAGG<br>GCCTCCAATGGGTGGCCAACA-TAAACCAAGATGGAACTGAGGA-ACACTATGTGGACTCTGTGCAGGGCCGCTTCACCATCTCCAGAGACAACGCCAGGAATT |
| 7 | IGHV3-48_* | TCTTTCGCACAGTGATAAACAGCCGTGTCCTCGGCTCTCAGGCTGAACAGTTGCAGAAACACTGAATTCCTGGCGTTGTCTCTGGAGATGGTGAAGCGGCCCTGCACAGAGTCCACATAGTGT-TCCTC<br>AGTTCCATCTTGGTTTA-TGTTGGCCACCCATTGGAGGCCCTTCCCTGGAGCCTGGCGGACCCAACTCATCCAAAAACTACTAAAACTGAATCCAGAGGCTGCAC |
| 2 | IGHV3-48_* | GAGGTGCAGGTGGTGGAGTCTGGGGAAGAGTTGCTACAGTTggggggggTCCCTGAGACTCTCCTGTGCAGCCTCTGGATTCACCCTCAGGCGAAACTGCATGAACTGAGTCCGCCAGGCTCCAGGGCAG<br>GGGCTGGAGTGGGTTTCAAACACTAGTGGAAGTTTTAATTTTAAAACCACATCTTAGGCAGACTCTCTGAAGGGCCGGTTCCCCATTTCCAGAGACAACGCCAA |
| 57 | IGHV3-48_04 | GAGGTGCAGGTGGTGGAGTCTGGGGAAGAGTTGCTACAGTTgggggggTCCCTGAGACTCTCCTGTGCAGCCTCTGGATTCACCCTCAGGCGAAACTGCATGAACTGAGTCCGCCAGGCTCCAGGGCAGG<br>GGCTGGAGTGGGTTTCAAACACTAGTGGAAGTTTTAATTTTAAAACCACATCTTAGGCAGACTCTCTGAAGGGCCGGTTCCCCATTTCCAGAGACAACGCCAAC |

| | | |
|---|---|---|
| 28 | IGHV3-48_* | TAAGACGCAGACGCGTCTCCGGCTCTCAGGCTGCTCATTCGCAGAAAGAGTGAGTTGTTGGCGTTGTCTCTGGAAATGGGGAACCGGCCCTTCAGAGAGTCTGC<br>CTAAGATGTGGTTTTAAAATTAAA<br>ACTTCCACTAGTGTTTGAAACCCACTCCAGCCCCTGCCCTGGAGCCTGGCGGACTCAGTTCATGCAGTTTCGCCTGAGGGTGAATCCAGAGGCTGCACAGGAGA<br>GT |
| 3 | IGHV3-48_* | TAAGACGCAGACGCGTCTCCGGCTCTCAGGCTGCTCATTCGCAGAAAGAGTGAGTTGTTGGCGTTGTCTCTGGAAATGGGGAACCGGCCCTTCAGAGAGTCTGC<br>CTAAGATGTGGTTTTAAAATTAAAA<br>CTTCCACTAGTGTTTGAAACCCACTCCAGCCCCTGCCCTGGAGCCTGGCGGACTCAGTTCATGCAGTTTCGCCTGAGGGTGAATCCAGAGGCTGCACAGGAGAG<br>T |
| 2 | IGHV3-72_* | ACAGTAATACACGGCTGTGTCCTCGGTTTTCAGGCTGTTCATTTGCAGATACAGCGTGTTTTTTGAATCATCTCTTGAGATGGTGAATCTGCCTTTCACGGGTGCA<br>GCGTAGTCTGTTGTCCCACCATCAGTTTTGCTTTTAATACGGCC-A---AC-CCA----C-T-CC---AG-CT--------CA-TCCA-G--G-<br>CGTTACTGAAAGTGAATCCAGAGGCTGCACAGGA |
| 2 | IGHV3-72_* | GAGGTGCAGCTGGTGGAGTCTGGGGGAGGCTTGGTAAAGCCTGGGGGGTCCCTTAGACTCTCCTGTGCAGCCTCTGGATTCACTTTCAGTAACG-C--C-TGGA-<br>TG--------AG-CT---GG-A-G----TGG-GT---T-GG<br>CCGTATTAAAAGCAAAACTGATGGTGGGACAACAGACTACGCTGCACCCGTGAAAGGCAGATTCACCATCTCAAGAGATGATTCAAAA |
| 2 | IGHV3-OR16-9_* | GAGGTGCAGTTGGGGGAGTCTGGGGGAGGCCTGGCACAGCCTGGAGGGTCCCTGAGATTCTCCTGTCCCTCCTCTCGATTCACACTT-<br>AATAATTTTAACATGGAG-TGGGTCCGCCAGGCTCCAGGGAAGG<br>GACTGGAGTGGGTTTCAGAGATTAGTG--AATAGC----A-AG-AACTTCGCAGACTCTGTGAAGGGCCGATTCAGCATCTCCAGAGACAACGCCAGGAGCT |
| 4 | IGHV4-28_* | CAGGTGCAGCTACAGGAGTCGGGCCCAGGACTGGCGAAGCCTTCGGAGACTTTGTCCCTCACCTGCAGTGTCTCTGGTGGCTCCAT--G-<br>AGTAATTACTTCTGGAGCTGGATCCGGCAGCCCGCC-GGAAGG<br>GACTGGAGTGGCTTGGGCGTATGTATACCAATGGGAGGACCGACTACAACCCCTCCCTCAAGAGTCGACTCACCATGTCAATAGACATGTCTAAGAACACC |
| 2 | IGHV4-28_* | CAGGTGCAGCTACAGGAGTCGGGCCCAGGACTGGCGAAGCCTTCGGAGACTTTGTCCCTCACCTGCAGTGTCTCTGGTGGCTCCAT--G-<br>AGTAATTACTTCTGGAGCTGGATCCGGCAGCCCGCC-GGAAGG<br>GACTGGAGTGGCTTGGGCGTATGTATACCAATGGGAGGACCGACTACAACCCCTCCCTCAAGAGTCGACTCACCATGTCAATAGACATGTCTAAGAACACC |
| 2 | IGHV4-34_* | CAGGTGCAGCTACAACAGTGGGGCGCAGGCCTGTTGAGGCCCTCGGAGACCCTGTCCCTCACCTGCGCTCTGTCTGGTGGGTCCTTCAGTGGTTACTACTGGAC<br>CTGGATTCGCCAGTCCCCAGGGAGG<br>GGACTGGAGTGGATTGGTGAAATCAATCATAGTGGGATTGGTGAAATCAATCATAGTGGGAGTACCAACTACAACCCGTCCCTCAAGAGTCGAGTCACCATAT<br>CA |
| 13 | IGHV4-39_* | CAACTGCAGCTGCAGGAGTCGGGCCCAGGACTGGTGAAGCCTTCGGAGACCCTGTCCCTCACCTGCGCTGTTGCTGGTGGCTCCATTAGCAGTGGCAGTTAGTT<br>CCGGGGCTGGGTCCGCCAGCCCCCAGGG<br>AAGGGGCTGGTGTGCATTGAGAATATGTACTAAACTGATATCACCCACTACAACCCGTCCCTCAGGAGTCGAGTCACCATATCCCAGTCACCATATCCGTAG |
| 2 | IGHV4-39_* | CTCGCGCACTAGTAAACAGCCGCGCCTGCGGCGGTCAGCGAGATCAGCGTCAGGGAGAGCTCGTTCTGCGTCGTGTCCTCGCACACGGTGACTCGACAC---A-<br>GGA-----TGGAGGAGGATG-TCCCACTATAG<br>GAGAGACTCTCAATCCACCCCAGCGCCATCCCTGGGGGCTGGCGGTTCCAGCCCCAGGACTCAATATTACATCTGATGGGGCCACCAGAGACAGTGCA |
| 8 | IGHV4-39_* | TGTCTCGCACAGTTAGAAAGAGCCGTGTCTGCGGCGGTCACAGACCTCAGCCTCAGGGAGAACCAGTTCTTGGACGGGTCTACGGATATGGTGACTGGGATATG<br>GTGACTCGACTCCTGAGGGACGGGTTGT<br>AGTGGGTGATATCAGTTTAGTACATATTCTCAATGCACACCAGCCCCTTCCCTGGGGGCTGGCGGACCCAGCCCCGGAACTAACTGCCACTGCTAATGGAGC |
| 2 | IGHV4-39_* | CCCTGTCCCTCACCTGCACTGTTTTTGGTGGCCCCATCGCCACTGACGA-<br>TCACCACTGGGGCTGGATCCGCCAGTCCCCGGGGAAGGGACTGGAGTGGATTTTGAGTGTCTATTATCGTGGGAGCGCCTATCG<br>TGGGAACACCTACTGCAATCCGTCCCTCAAGGGACGAGTCGCCGTGTCCGTAGAGGCGTCCAGGAACAACTTCTCCCTGAAACTGAGCTCTGTGACCGCC |
| 6 | IGHV4-39_* | TCTCGCACAGTAATAGACAGCCGTGTCTGCGGCGGTCACAGAGCTCAGTTTCAGGGAGAAGTTGTTCCTGGACGCCTCTACGGACACGGCGACTCGTCCCTTGA<br>GGGACGGATTGCAGTAGGTGTTCCCACGA<br>TAGGCGCTCCCACGATAATAGACACTCAAAATCCACTCCAGTCCCTTCCCCGGGGACTGGCGGATCCAGCCCCAGTGGTGATCGTC-AGTGGCGATGGGGC |
| 12 | IGHV4-39_* | TGCTGCTGCAGGAGTCGGGCCCAGGACTGGTGAAGCCTTCGGAGACCCTGTCCCTCACCTGCACTGTTTTTGGTGGCCCCATCGCCACT-<br>GACGATCACCACTGGGGCTGGATCCGCCAGTCCCCGGGGAAGGG<br>ACTGGAGTGGATTTTGAGTGTCTATTATCGTGGGAGCGCCTATCGTGGGAACACCTACTGCAATCCGTCCCTCAAGGGACGAGTCGCC<br>GTGTCCGTAGAG |
| 2 | IGHV4-OR15-8_* | CAGGTGCAGTTGCAGGAGTCGGGCCCAGGGCTGGTGAAGGCTTCGGAGACCCTGTCCCTCATGTGCACTGTCTCTGGTGACCTCACCAACACTTAT-<br>ACTACTGGAACTGGATCCGGCAG-GCCCGGGGAGG<br>GGACTCGAATGGATTGGGTTTGTC-ACCAACAGTGGGAGAAGCAACTCCAACCCCT-CCTCAAGAGTCGAGTCACCATATCAATAGACACATCCAAGAACCA |
| 2 | IGHV4-OR15-8_* | TCTCTCGCACAGTAATAAACGGCCGTGTCCGCAGAGGTCACAGAGGTCAGCCTCAGGGAGAACTGGTTCTTGGATGTGTCTATTGATATGGTGACTCGACTCTT<br>GAAGGAGGGGTTGGAGTTGCTTCTCCCACT<br>GTTGGT-GACAAACCCAATCCATTCGAGTCCCCTCCCCGGGGCCTGCCGGATCCAGTTCCAGTAGTAAGTGTTGGTGA-GGT--CACCAGAGACAGTGCA |
| 4 | IGLV2-23_* | CAGTCTGCCCTGACTCAGCCTGCCTCCGTGTCTGGGTTTCCTGGACAGTCGAACACCATCCCCTGCACTGGAGCCAGCAGGGATGTTGGGGGATATTAATTTCTC<br>GCCTGGCTGCAACACCACCCAGACAAGGT<br>CCCCAAACTCACGATTGATGACGTCTCTAAGCGGCCCGCAGATACTACTACCTGCTTTTCTGGCTCCAAGTCTGGCAACGCGCCCTCCCTGACAGTCGCT |
| 9 | IGLV2-23_* | GCAGAAG-AA<br>AATCAGCCTCGTCCCCAGCCGGGAGCCCAGCGACTGTCAGGGAGGGCGCGTTGCCAGACTTGGAGCCAGAAAAGCAGGTAGTAGTATCTGCGGGCCGCTTAGA<br>GACGTCATCAATCGTGAGTTTGGGGACCTT<br>GTCTGGGTGGTGTTGCAGCCAGGCGAGAAATTAATATCCCCCAACATCCCTGCTGGCTCCAGTGCAGGGGATGGTGTTCGACTGTCCAGG |
| 2 | IGLV3-16_* | TCCTATGAGCTGACACAGCCACCCTCGGTGTCAGTGTCGCTAGGACAGATGGCCAGGATCGCCTGCTCCGGGGGAACCATCGTCAGAAAAATGTGATTCTTTCT<br>TGTACCAG-A-AATT-AGGCCAGTTCCCTGTG<br>CTGGT----------------GAGAGGCCCTCAGGGATCCCTGACCGATTCTCTGGCTCCACTTCAGGGACAATAGTCACATTGACCAGCAATGGAGT |

* Indicate events belong to this gene subclass but unknown annotation.
